# Age and diet modulate the insulin-sensitizing effects of exercise: a tracer-based oral glucose tolerance test

**DOI:** 10.1101/2023.03.18.533083

**Authors:** Marcel A. Vieira-Lara, Aaffien C. Reijne, Serj Koshian, Jolita Ciapaite, Fentaw Abegaz, Alzbeta Talarovicova, Theo H. van Dijk, Christian J. Versloot, Robert H.J. Bandsma, Justina C. Wolters, Albert K. Groen, Dirk-Jan Reijngoud, Gertjan van Dijk, Barbara M. Bakker

**Author notes:** These authors contributed equally (Marcel A. Vieira-Lara, Aaffien C. Reijne).

## Abstract

Diet modulates the development of insulin resistance during aging. This includes tissue-specific alterations in insulin signaling and mitochondrial function, which ultimately affect glucose homeostasis. Exercise stimulates glucose clearance, mitochondrial lipid oxidation and enhances insulin sensitivity. It is not well known how exercise interacts with age and diet in the development of insulin resistance. To investigate this, oral glucose tolerance tests (OGTT) with a tracer were conducted in mice ranging from 4 to 21 months of age, fed a low- (LFD) or high-fat diet (HFD), with or without life-long voluntary access to a running wheel (RW). We developed a computational model to derive glucose fluxes, which were commensurate with independent values from steady-state tracer infusions. Both insulin sensitivity indices derived for peripheral tissues and liver (IS-P and IS-L, respectively) were steeply decreased by aging and a HFD. This preceded the age-dependent decline in the mitochondrial capacity to oxidize lipids. In LFD young animals, RW access enhanced the IS-P concomitantly with the muscle β-oxidation capacity. Surprisingly, RW access completely prevented the age-dependent IS-L decrease, but only in LFD animals. This study indicates, therefore, that endurance exercise can improve the age-dependent decline in organ-specific IS mostly in the context of a healthy diet.

## Introduction

Aging, diet and exercise have a profound effect on insulin sensitivity and glucose homeostasis (1–3). As a consequence, they affect the emergence of insulin resistance, a pathophysiological process that precedes the development of type 2 diabetes. Insulin resistance is characterized by decreased insulin-stimulated glucose uptake and decreased insulin-dependent suppression of the endogenous glucose production (EGP) (2). In the postprandial state, the skeletal muscle represents one of the major sites for glucose clearance together with the central nervous system, whereas the main site of EGP is the liver (4–6). Insulin resistance in the skeletal muscle and the liver can be induced by the accumulation of lipids, which involves the inhibition of the insulin signaling cascade by complex lipids (2). This lipid-induced insulin resistance can be elicited in mice fed a high-fat diet (7). It is questioned, however, whether insulin resistance development is caused by an overload of substrate (8) or by a loss of mitochondrial flexibility and/or function (9; 10), or a combination of both.

Aging exacerbates the susceptibility to lipid-induced insulin resistance (11; 12), which amongst other factors involves changes in lipid handling, both in the skeletal muscle (11; 13) and the liver (14). The main route for lipid oxidation is the mitochondrial fatty-acid β-oxidation, with its flux being mostly controlled at the level of carnitine palmitoyltransferase 1 (CPT1) (13; 15). We previously reported that an inability of old animals to upregulate muscle CPT1B (muscle isoform) in response to a high-fat diet functioned as a basis for insulin resistance development (13). Aging is also a risk factor for insulin resistance on its own (1), and its association with sarcopenia (loss of muscle mass) (16) may further exacerbate insulin resistance (17).

In humans, both resistance (strength) and endurance (aerobic) exercise have been associated with improved insulin sensitivity (18; 19). Whereas resistance training rescues age-related muscle loss (20), endurance training is generally more effective in improving mitochondrial function in aging (16; 21; 22). The high mitochondrial capacity of endurance-trained athletes was shown to protect against lipid-induced insulin resistance (23; 24). With aging, however, the link between mitochondrial capacity and insulin sensitivity as a result of exercise may disappear: it was found that aerobic exercise improved oxidative capacity irrespective of age, whereas it improved insulin sensitivity only in young subjects (24). Access to a running wheel (RW) is a widely used protocol to study the effects of endurance exercise in mice (25). In line with the above, in mice ranging from 6 to 24 months old, RW access increased mitochondrial content and oxidative capacity in the quadriceps, irrespective of whether a low- or high-fat diet was given. This was dissociated from the age-related decline in muscle weight (26). Finally, exercise does not only affect insulin sensitivity and glucose handling in the muscle, but also in the liver (27; 28). After long-term endurance training, the ability of the liver to take up glucose increases and so does its sensitivity to insulin (27).

Although the effects of endurance exercise have been studied at different ages and on different diets, it has not yet been systematically addressed how these three factors interact in the development of muscle and liver insulin resistance. Moreover, a tissue-specific analysis of insulin resistance and glucose handling is crucial to dissect the role of mitochondrial function in insulin resistance. For this purpose, indices that distinguish muscle (peripheral) and liver insulin sensitivity are necessary. The frequently used HOMA-IR index (Homeostasis Model Assessment of Insulin Resistance) is a measure of whole-body insulin sensitivity, based on fasting glucose levels. An Oral Glucose Tolerance Test (OGTT) is used to derive the Muscle Insulin Sensitivity Index (MISI, focused on peripheral insulin sensitivity) or a Matsuda index (whole-body insulin sensitivity) during a glucose load (5; 29; 30). Including a stable-isotope labeled glucose tracer to either oral or intravenous glucose tolerance tests, allows to (i) dissect the contribution of the EGP from that of the glucose elimination by the muscle and other peripheral tissues and (ii) quantify insulin sensitivity in a tissue-specific manner (31–34). In the present study, we combine this advantage of a tracer with the OGTT.

We here investigate the combined effects of age, diet and voluntary wheel running on glucose homeostasis and tissue-specific insulin sensitivity in a cohort of aging mice about which we reported previously (26; 35). To this end, we applied an existing OGTT protocol with [6,6-^2^H_2_]-glucose in mice (36) and adapted a computational model from earlier work in humans (37).

## Research Design and Methods

### Mouse experiments and ethics approval

Mouse experiments were performed as previously described (26; 35). Briefly, male C57BL/6JOlaHsd mice (Jackson Laboratory, Bar Harbor, ME, USA) were maintained at 22 °C on a 12h/12h light/dark cycle and *ad libitum* access to a low-fat diet (LFD, 6% calories from fat, 1% w/w sugar; AMII 2141, HopeFarms BV, Woerden, NL). On postnatal day 28, mice were placed in individual cages (Makrolon type II, Bayer, Germany) and given either the same LFD or a high-fat high-sucrose diet during their entire lifetime (HFD, 45% calories from fat, 20% w/w sucrose; 4031.09, HopeFarms BV, Woerden, NL). A total of 128 mice were randomly subdivided into voluntary running-wheel groups (RW) or sedentary controls (Ctrl). At the age of 4, 9, 15 or 21 months (used throughout the text to define the age groups), animals underwent an OGTT with tracer (Fig. 1A). The same animals underwent steady-state intravenous infusion experiments (38) at the age of 6, 12, 18 or 24 months, respectively, after which animals were terminated and the quadriceps and liver were collected, as previously described (26).

**Figure 1:**
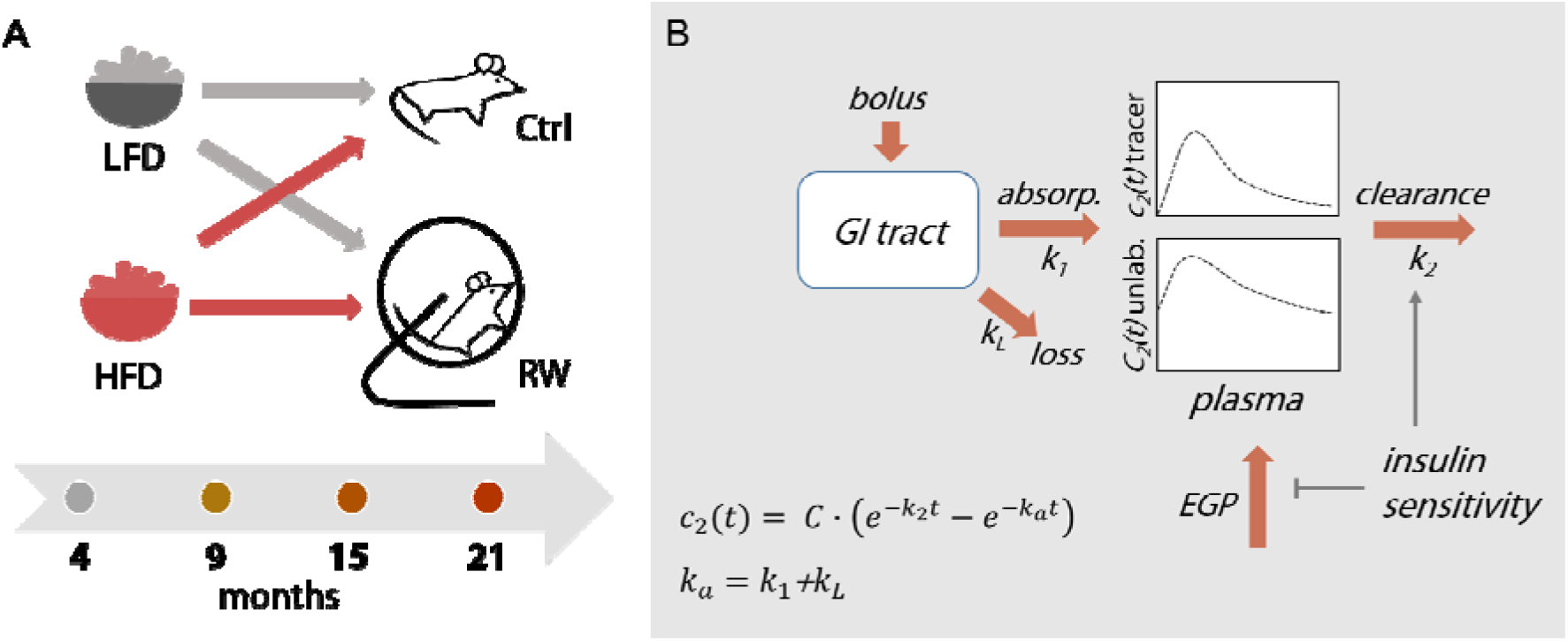
Schematic representation of experimental design (A) and compartment model for data analysis (B). LFD: low-fat diet, HFD: high-fat high-sucrose diet, Ctrl: sedentary mice, RW: mice with voluntary access to a running wheel. GI tract: gastrointestinal tract; absorp.: absorption flux into the plasma compartment, represented by the rate constant *k_1_*; loss: loss flux from the GI compartment, represented by the rate constant *k_L_*. Both tracer and unlabeled glucose in the plasma compartment can be cleared from the circulation, represented by the rate constant *k_2_*. The endogenous glucose production (EGP) by the liver feeds into the unlabeled glucose pool in the plasma. Insulin enhances the clearance of both tracer and glucose and inhibits the EGP. The apparent absorption constant *k_a_* obtained from the tracer curves equals *k_1_* + *k_L_* (See Materials and Methods for details).

To determine the bioavailability of the tracer, independent experiments were conducted in 3-week-old male C57BL/6J mice (Jackson Laboratory, Bar Harbor, ME, USA), fed a LFD (Teklab Custom Diet TD.10098, Envigo Teklad Diets, Madison, WI, USA) for 2 weeks prior to tracer measurements. Animal experiments were approved by the Dutch Central Authority for Scientific Procedures on Animals (CCD) and by the University of Groningen Ethical Committee for Animal Experiments.

### Oral Glucose Tolerance Test with glucose tracer (tracer OGTT)

On the day of the experiment, food was removed in the morning at 7 h (winter) or 8 h (summer time). The OGTT was conducted after a fasting period of 6 h, i.e. at 13 h or 14 h. Body weight was measured before the start of the experiment. Access to a RW was maintained during the OGTT timeframe. In the RW groups, the last bout of intense exercise occurred 6-7 h before the ingestion of the glucose bolus, according to data on average RW revolutions at different ages (Fig. S1). At time point zero a glucose bolus of 1 g·kg^-1^ (5.5·10^3^ μmol·kg^-1^) was administered orally. It consisted of 0.7 g·kg^-1^ (3.9·10^3^ μmol·kg^-1^) unlabeled glucose and 0.3 g·kg^-1^ (1.6·10^3^ μmol·kg^-1^) [6,6-^2^H_2_]-glucose (tracer). Total blood glucose (Lifescan Euroflash; Lifescan Benelux, Beerse, Belgium) was measured from the tail vein, from which bloodspots were also collected. Insulin was measured according to manufacturer instructions (Ultrasensitive Mouse Insulin kit, Mercodia, Uppsala, Sweden). The fractional contribution of the administered tracer in blood glucose (described below) was measured in air-dried blood spots. Glucose and tracer enrichment was measured at 0, 15, 30, 45, 60, 75, 90, 105 and 120 min and insulin was measured at 0, 30 and 60 min.

### Bioavailability

Five-week-old animals from an independent cohort were fasted for 4h and received an intravenous bolus of [6,6-^2^H_2_]-glucose (1.0·10^3^ μmol·kg^-1^) injected retro-orbitally. This was directly followed by an oral bolus containing glucose (2.0·10^3^ μmol·kg^-1^) and [U-^13^C]-glucose (70 μmol·kg^-1^). Bloodspots were collected at 0, 10, 20, 30, 40, 50, 60 and 90 min and glucose was measured as described above. Air-dried bloodspots were used for posterior analysis of fractional abundance.

### Gas chromatography coupled to mass spectrometry (GC-MS) and isotope correction

Fractional distributions of [6,6-^2^H_2_]-glucose and [U-^13^C]-glucose in bloodspots were measured according to van Dijk et al. (38). Glucose was converted into its pentaacetate derivative by adding pyridine/acetic anhydride mixture. All samples were analyzed by gas chromatography coupled to mass spectrometry (Agilent 9575C inert MSD; Agilent Technologies, Amstelveen, The Netherlands). Derivatives were separated on AT-1701 30 m x 0.25 mm ID (0.25 μm film thickness) capillary column (Alltech, Breda, The Netherlands).

The monitored isotopologue distribution (*m/z* 408-414) was corrected for the natural abundance of isotopes based on the approach using the measured isotopologue distribution in the baseline samples, as described in the Supp. Material S1. Tracer concentrations were obtained by multiplying the corrected fractional abundance of the *m/z* 410 isotopologue for [6,6-^2^H_2_]-glucose and *m/z* 414 for [U-^13^C]-glucose by the corresponding total glucose concentration. Unlabeled glucose concentrations were obtained by subtracting tracer concentrations from the total measured glucose concentrations.

### Modeling strategy

To derive the kinetic constants from the tracer data, we considered two compartments: the gastrointestinal tract compartment (compartment 1) and the blood plasma (compartment 2) (Fig. 1B). The pool sizes (*q*) and concentrations (*c*) of the tracer in these compartments are denoted by *q_1_, q_2_, c_1_* and *c_2_*, and those of unlabeled glucose analogously by *Q_1_, Q_2_, C_1_* and *C_2_*, with the subscript specifying the compartment. Pool sizes *Q* and *q* were expressed in μmol·kg^-1^ (referring to kg body weight), while concentrations *C* and *c* were expressed in mM. Inspired by the experimental setup, the model was confined to the *pool size* of the glucose that was administered in the stomach by oral gavage and dosed in μmol·kg^-1^ (*Q_1_*, *q_1_*), and the plasma *concentrations* (*C_2_, c_2_*) that were measured in mM. All modeling details can be found in Appendix 1.

### CPT1 quantification and oxidative capacity

Mitochondria were isolated from quadriceps and liver tissue, as previously described (26; 39). CPT1A and CPT1B protein levels were quantified by targeted proteomics in isolated mitochondrial suspensions (40). Oxygen consumption rates were measured in MiR05 buffer (41) in a two-channel high-resolution Oroboros oxygraph-2k (Oroboros, Innsbruck, Austria). Palmitoyl-CoA (25 μM), L-carnitine (2 mM) and malate (2 mM) were used as substrates. Maximal ADP-stimulated respiration was measured following the addition of 1.5 U/mL hexokinase, 12.5 mM glucose and 1 mM ATP. Protein content was measured with the Pierce BCA Protein Assay Kit (Thermo Fisher 23225). The ratio of cytosolic to mitochondrial protein was determined as previously described (26) and used to express CPT1 content and oxidative capacity per total tissue protein.

### Software and analysis

Correction for natural abundance was performed in Excel 2019. All other calculations and simulations were conducted in Python (v 3.7), using Jupyter Notebook (v 5.7.4). Parameters from the tracer kinetics and curves for the unlabeled glucose were estimated with the *minimize* function from the *lmfit* Python package (Levenberg-Marquardt method, non-linear least squares). Data visualization and statistical analysis were conducted either in Python (*matplotlib, scipy*) or GraphPad Prism (v9.0, GraphPad Software, San Diego, California USA). Statistical analyzes comparing the effects of age, diet and exercise, as well as their interaction, were conducted by three-way ANOVA.

### Data and Resource

All data generated or analyzed during this study are included in the published article (and its online supplementary files). The resources generated during the current study can be found at github.com/mvieiralara/OGTT_modelling.

## Results

### HFD reduces glucose elimination in an age-dependent manner

Mice fed either a low-fat or high-fat high-sucrose diet (LFD and HFD, respectively) were divided into a sedentary group (Ctrl) and a group with voluntary access to a running wheel (RW). Physiological characteristics of these mice have been previously described elsewhere (35). To study glucose kinetics and insulin sensitivity, a tracer OGTT was conducted in mice that were 4, 9, 15 or 21 months old (Fig. 1A). A computational model consisting of glucose absorption, loss, clearance and EGP was built (Fig. 1B and Appendix 1). The tracer time course data were fitted to the equation in Fig. 1B (Fig. 2 and Supp. Material 3).

**Figure 2:**
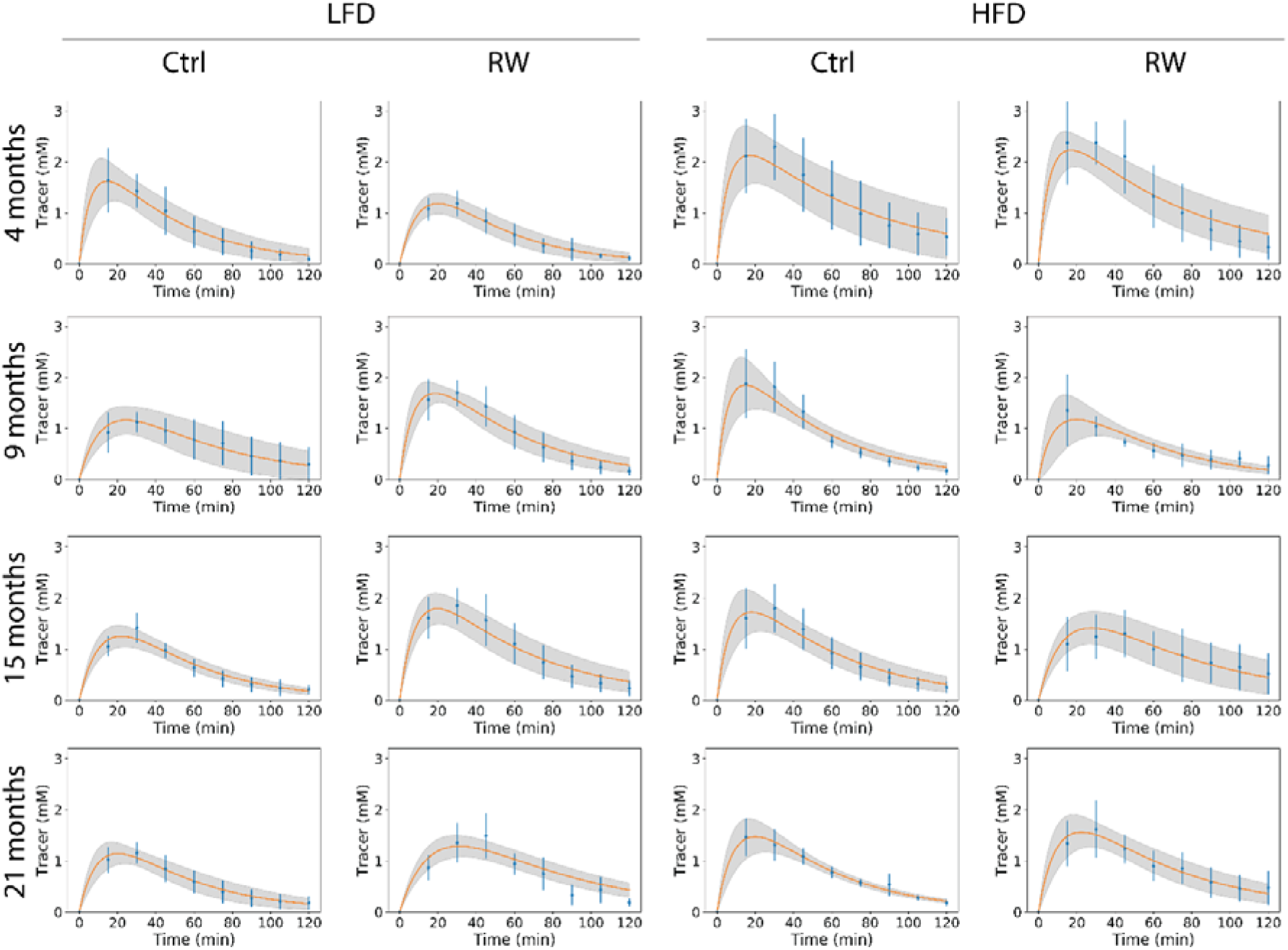
Curve fits for time courses of the [6,6-^2^H_2_]-glucose tracer during OGTT. Each column represents a different diet and activity group, whereas each row represents a different age. LFD: low-fat diet, HFD: high-fat high-sucrose diet, Ctrl: sedentary mice, RW: mice with access to a running wheel. Means ± SD for each time point are shown in blue. The average curve fit from all animals per experimental group (*c_2_* in the model) is shown in orange, with the SD of the fitted curves represented by the shaded area. N=6-8 per group.

The apparent rate constant of absorbance *k_a_* (Fig. 1B) was higher in the HFD than in the LFD groups (pdiet < 0.01, Fig. 3A) and decreased with advanced age (page <0.001, Fig. 3A). The apparent rate constant *k_2_* represents the fractional clearance rate of glucose from the plasma compartment (‘glucose effectiveness’). The HFD decreased *k_2_* (vs LFD, p_diet_ < 0.001), while *k_2_* responded differently to aging in each diet group (p_age_×_d_i_et_<0.001), with no effect of RW access. In an independent tracer experiment in which intravenous (IV) and oral administration of the tracer were compared, the bioavailability *F* was determined to be 0.8 (Fig. 3C-F), implying that 80% of the tracer is absorbed and 20% is lost from our observation. Thus, *k_1_* = 0.8·*k_a_* and *k_L_* = 0.2·*k_a_* (cf. Fig. 1B). An 80% bioavailability is in agreement with previous studies in humans and dogs (42; 43). The apparent volume of distribution (*Vol*, eq. 14) was lower in the HFD group than in the LFD group (p_diet_ <0.001), and responded in a diet-dependent manner to the RW (p_diet×RW_ < 0.01) and age (p_diet×age_ < 0.05) (Fig. 3G). In summary, all kinetic constants in the model could be obtained from the tracer OGTT, if the bioavailability is independently determined. Moreover, the rate constant of glucose clearance was reduced by the HFD in an age-dependent manner.

**Figure 3:**
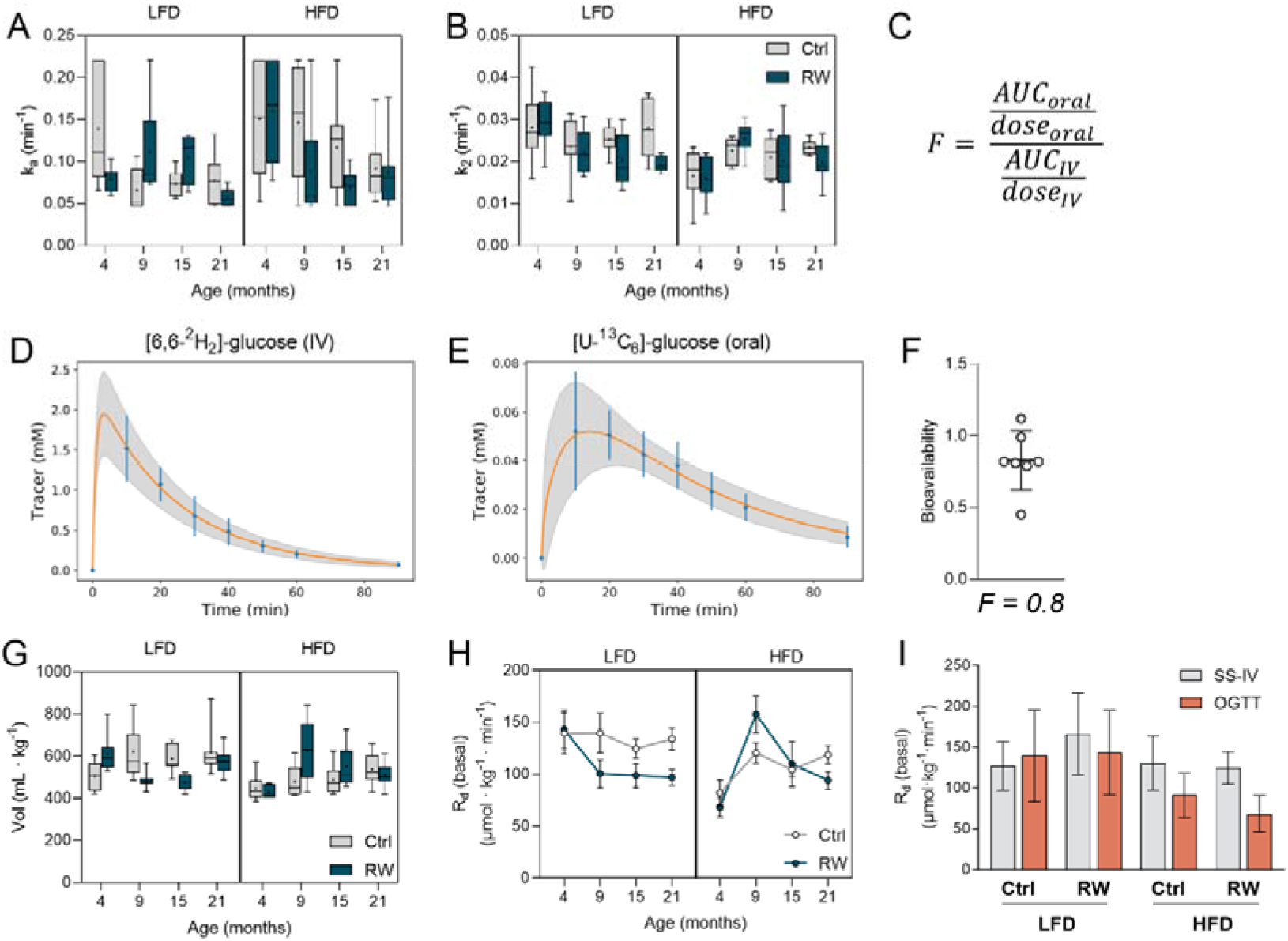
The HFD reduced both the rate constant and the rate of glucose elimination. Kinetic constants *k_a_* (A) and *k_2_* (B) calculated from curve fits. Equation for calculating the bioavailability *F* based on IV and oral experiments, in which the areas under the curve (AUC) are corrected for the dose and then divided (C). To calculate *F*, curves were fitted to tracer time courses following either IV (D) or oral administration (E) of glucose to 5-week-old mice. Data points are shown in blue as mean ± SD (n=8) and the average curve fit from all animals in orange, with the SD of the fitted curve is represented by the shaded area. Calculated bioavailability *F* (F). Apparent volume of distribution (*Vol*) assuming a constant *F* for all groups (G). Basal rate of disappearance of glucose (*Rd*) calculated from the OGTT data (mean ± SEM) (H). Comparison between *R_d_* calculated from the tracer OGTT in 4-month-old animals and steady-state intravenous infusion experiments conducted in the same animals at 6 months. Data are shown as mean ± SD, n=6-8 (I). LFD: low-fat diet, HFD: high-fat high-sucrose diet, Ctrl: sedentary mice, RW: mice with voluntary access to a running wheel.

The HFD treatment reduced the basal glucose disappearance rate (*R_d_*) (p_diet_ < 0.05), while *R_d_* was affected by age in a diet-dependent manner (p_age×diet_ < 0.001) (Fig. 3H). Subsequently, these outcomes were compared to those of classical steady-state intravenous infusion (SS-IV) experiments (38), which allow for a model-independent measurement of basal glucose fluxes. In the 4 month-old group, basal 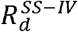 values (eq. 26) were in the same range as those obtained from the tracer OGTT (Fig. 3I). Although this was true for all different ages, the biological trends were not the same (Fig. S2). This reflects the nature of the experiment: in the OGTT the *k_2_* was estimated over the whole time course, during which the glucose bolus stimulated the release of hormones such as insulin. In the steady-state experiment, only tracer was infused, which should have barely affected hormone production.

### Advanced age leads to decreased endogenous glucose production

The time courses of unlabeled glucose (Fig. 4) depend on the same kinetic parameters as that of the tracer and, in addition, on the EGP (Fig. 1B). The variation within the groups was larger for unlabeled than for labeled glucose (cf. Fig. 2 and 4), indicating more variation in the EGP component, which may reflect stress due to animal handling (44). Individual curves that did not show the typical absorption followed by a clearance phase, occurred in all experimental groups and were excluded (Fig. S3). Fasting glucose values (Fig. S4) were elevated in the HFD group (p_diet_ < 0.01) and reduced by advanced aging (p_age_ < 0.05), which in turn was modulated by the RW (p_RW×age_ < 0.05).

**Figure 4:**
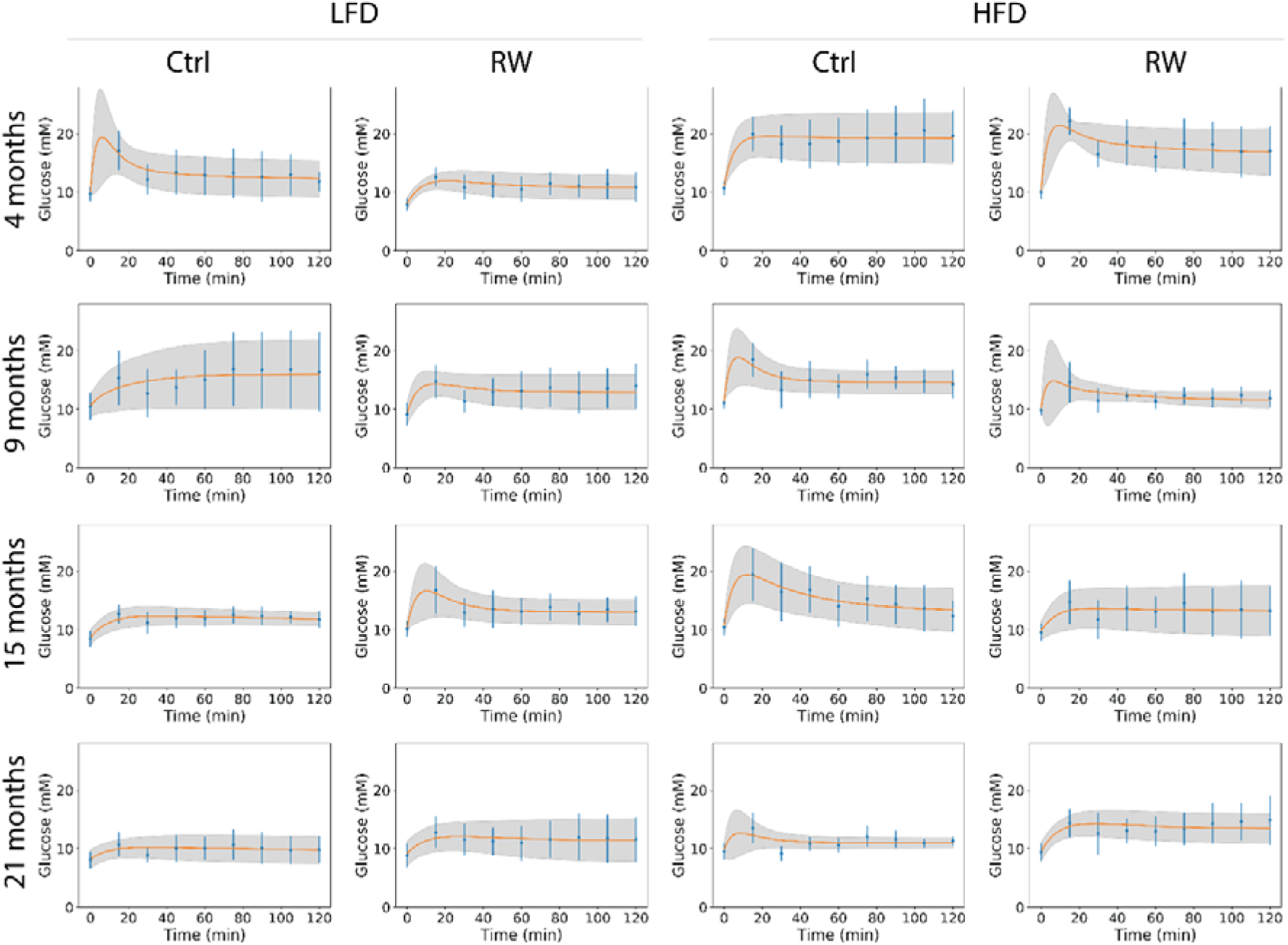
Curve fits of unlabeled glucose time courses during OGTT. Each column represents a different diet and activity group, whereas each row represents a different age. LFD: low-fat diet, HFD: high-fat high-sucrose diet, Ctrl: sedentary mice, RW: mice with voluntary access to a running wheel. Means ± SD for each time point are shown in blue. The average curve fit (*C_2_* in the model) is shown in orange, with the SD represented by the shaded area. N=6-8 per group.

After administration of the glucose bolus, the EGP decreased initially in most groups, reaching a minimum between 5 and 20 minutes, before reaching a new stationary level (Fig. 5A). The initial decline can be attributed to the inhibitory effect of increased plasma insulin and glucose (45). The steady-state EGP was calculated as the average of the last 30 minutes (Fig. 5B), and the overall average EGP was calculated over the entire period from 5 to 120 min (Fig. 5C). Higher age decreased both steady-state and average EGP (p_age_ < 0.01) (Fig. 5B-C), which is in line with the usually lower glucose levels in older animals (Fig. 4). The RW generally decreased the steady-state and average EGP (vs Ctrl, p_RW_ < 0.05), mostly in the LFD group (pRW×diet < 0.05). The calculation of EGP in μmol·kg^-1^·min^-1^ (Fig. 5) depends on the assumption of constant bioavailability. When expressed in mM·min^-1^, however, the EGP* was independent of the bioavailability (Appendix 1) and followed the same trend (Fig. S5A) as in Fig. 5A. Similar patterns were also observed for steady-state and average EGP* (Fig. S5B-C). This highlights that the differences between groups are independent of the assumption of a constant bioavailability.

**Figure 5:**
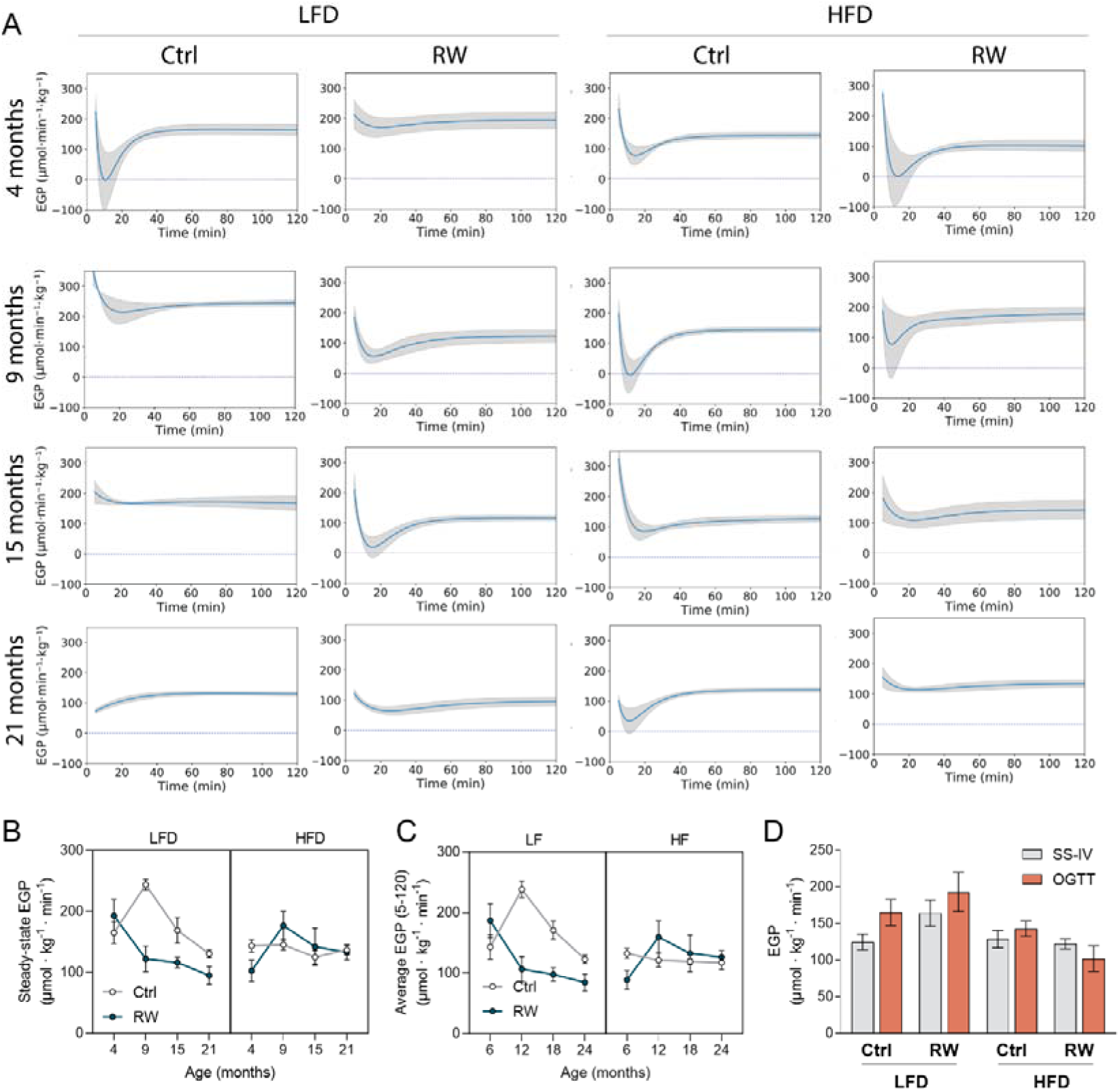
EGP is reduced by aging in the LFD group. Time courses for EGP (A). Each column represents a different diet and activity group, whereas each row represents a different age. LFD: low-fat diet, HFD: high-fat high-sucrose diet, Ctrl: sedentary mice, RW: mice with voluntary access to a running wheel. Mean EGP (line) ± SEM (shaded area) is shown per experimental group. Steady-state EGP values (B) calculated from the curves (mean ± SEM). Time-averaged EGP values (C) obtained from OGTT timeframe (5-120 min, mean ± SEM). Comparison between steady-state EGP values (mean ± SD) obtained from the tracer OGTT in 4-month-old animals and steady-state intravenous infusion experiments conducted in the same animals at 6 months (D). N=2-8 per group.

The steady-state EGP derived from the tracer OGTT (EGP^OGTT^) was compared with independent steady-state intravenous experiments conducted in the same cohort of animals. This yielded a model-independent basal EGP (EGP^SS-IV^). EGP^OGTT^ and EGP^SS-IV^ were similar to each other in 4 month-old animals (Fig. 5D). Although the same trend was observed in all age groups, the tracer OGTT usually yielded higher EGP values than the steady-state experiment (Fig. 5D and S5D).

### RW mostly affects liver, and not muscle, insulin sensitivity

Both age and HFD strongly increased the insulin levels during the OGTT (p_age_ and p_diet_ < 0.001), in line with previous results (13; 35), while RW access decreased insulin levels by around 20% (p_RW_ < 0.01) (Fig. 6A). HOMA-IR, a classical surrogate index for whole-body insulin resistance, followed a similar pattern as the average insulin levels (Fig. 6B). Age and HFD (p_age_, p_diet_ < 0.001) increased HOMA-IR, whereas the RW reduced it (p_RW_ < 0.05). The effect of the RW was particularly pronounced in the LFD group.

**Figure 6:**
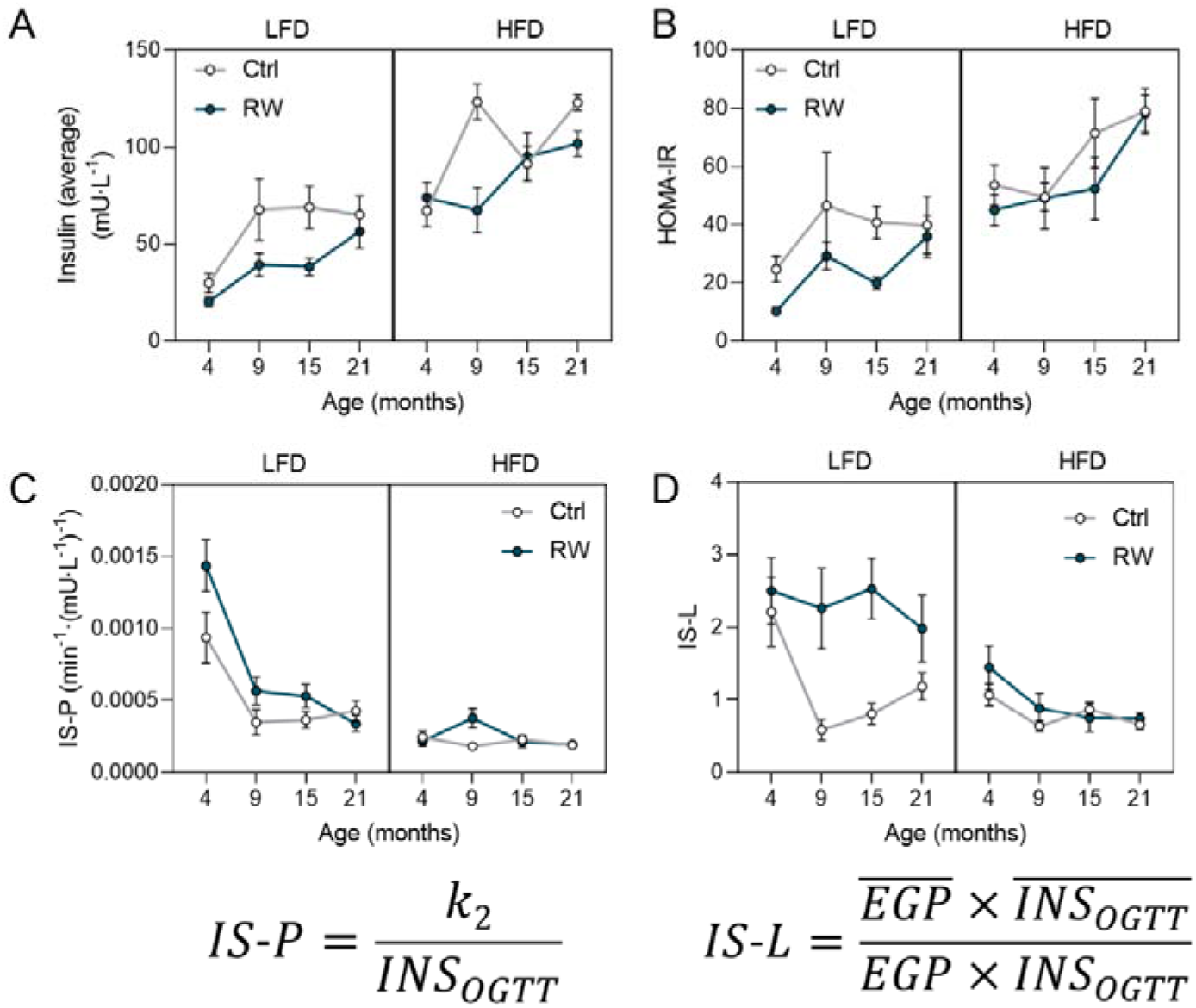
Peripheral and central insulin sensitivity are regulated by age, diet and RW. Average insulin levels during the OGTT (0, 30 and 60 min) (A). HOMA-IR calculated from fasting glucose and fasting insulin levels (B). IS-P (C) and IS-L (D) for tracer OGTT, calculated according to the featured equations. *INS_OGTT_* is the time-average insulin levels per animal, *EGP* is the time-average EGP per animal (5-120 min). 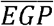 and 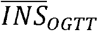 are the average of *EGP* and *INS_OGTT_* values, respectively, for all 16 experimental groups. LFD: low-fat diet, HFD: high-fat high-sucrose diet, Ctrl: sedentary mice, RW: mice with voluntary access to a running wheel. Data are shown as mean ± SEM, n=2-8 per group.

Despite its wide application, the HOMA-IR fails to capture organ-specific insulin resistance (5; 46). We defined a novel insulin sensitivity index for peripheral tissues (IS-P) as the fractional clearance (*k_2_*, min^-1^) divided by the insulin concentration (mU·L^-1^) (Eq. 29 Appendix 1, Fig. 6). In the LFD groups, the IS-P decreased strongly between 4 and 9 months, in sedentary as well as RW groups. In the HFD groups the IS-P was already low at a young age (4 months) and did not decline further during aging (Fig. 6C). Together, age, diet and their interaction explained more than 50% of the variation in the dataset (p_age_, p_diet_, p_age×diet_ < 0.001). RW access significantly increased the IS-P (p_RW_ < 0.05), particularly in the 4 months old LFD group (around 50% increase vs Ctrl) (Fig. 6C). RW access alone, however, explained less than 2% of the variation in the data. The interaction terms RW×diet and RW×age showed p-values of 0.06 and 0.07, respectively, which points to a trend of differential effects of the RW in aged and HFD-fed groups.

Insulin also regulates the EGP, although this regulation is more complex to analyze (47). We adapted a hepatic insulin sensitivity index which is originally based on fasting EGP and insulin levels (29). Instead, we used the time-averaged EGP and insulin levels to apply the index to the postprandial state mimicked by the tracer OGTT (IS-L, eq. 30 Appendix 1 and Fig. 6). Similar to the IS-P, the insulin sensitivity index for the liver (IS-L) was strongly reduced by aging, and again to a smaller extent in the HFD group where the IS-L was already low already from 4 months of age (Fig. 6D). Surprisingly, in the LFD group RW access strongly enhanced the IS-L, even much more than it affected the IS-P, thus delaying the age-related loss of the hepatic insulin sensitivity (Fig. 6D). Using the EGP* in mM·min^-1^ (Fig. S5C) yielded the same pattern for the IS-L (Fig. S6A), which reiterates that the effects of age, diet and RW do not depend on the assumption of constant tracer bioavailability.

Both IS-P and IS-L correlated with the HOMA-IR (Fig. S6B-C). The differences between IS-P and IS-L curves warrant a strict dissection of whole-body insulin sensitivity by its peripheral and liver components.

### RW access increases the β-oxidation capacity primarily in the muscle of LFD-fed animals

The quantitative dissection of peripheral and liver insulin sensitivity allowed us to relate these to mitochondrial function in specific tissues. Alterations in mitochondrial fatty-acid β-oxidation can modulate lipid clearance and impact insulin resistance development (48; 49). The content of CPT1B in the skeletal muscle declined with age (p_age_ < 0.001) in all analyzed groups (Fig. 7A), which aligns with an independent study (13). The HFD strongly upregulated muscle CPT1B content (p_diet_ < 0.001), whereas RW access increased it only in LFD animals (p_RW_ < 0.01, p_RW×diet_ < 0.001). In the skeletal muscle, the measured oxidative capacity with palmitoyl-CoA as substrate roughly correlated with the CPT1B levels (Fig. 7C, S7A). However, LFD Ctrl mice did not show a decline in oxidative capacity with age, in contrast to the CPT1B level (cf. Fig. 7A and C). Of note, in the youngest and oldest groups, both CPT1B content and oxidative capacity followed the same pattern of age-dependent loss of flexibility to the HFD, as previously reported by us (Fig. S7C-D).

**Figure 7:**
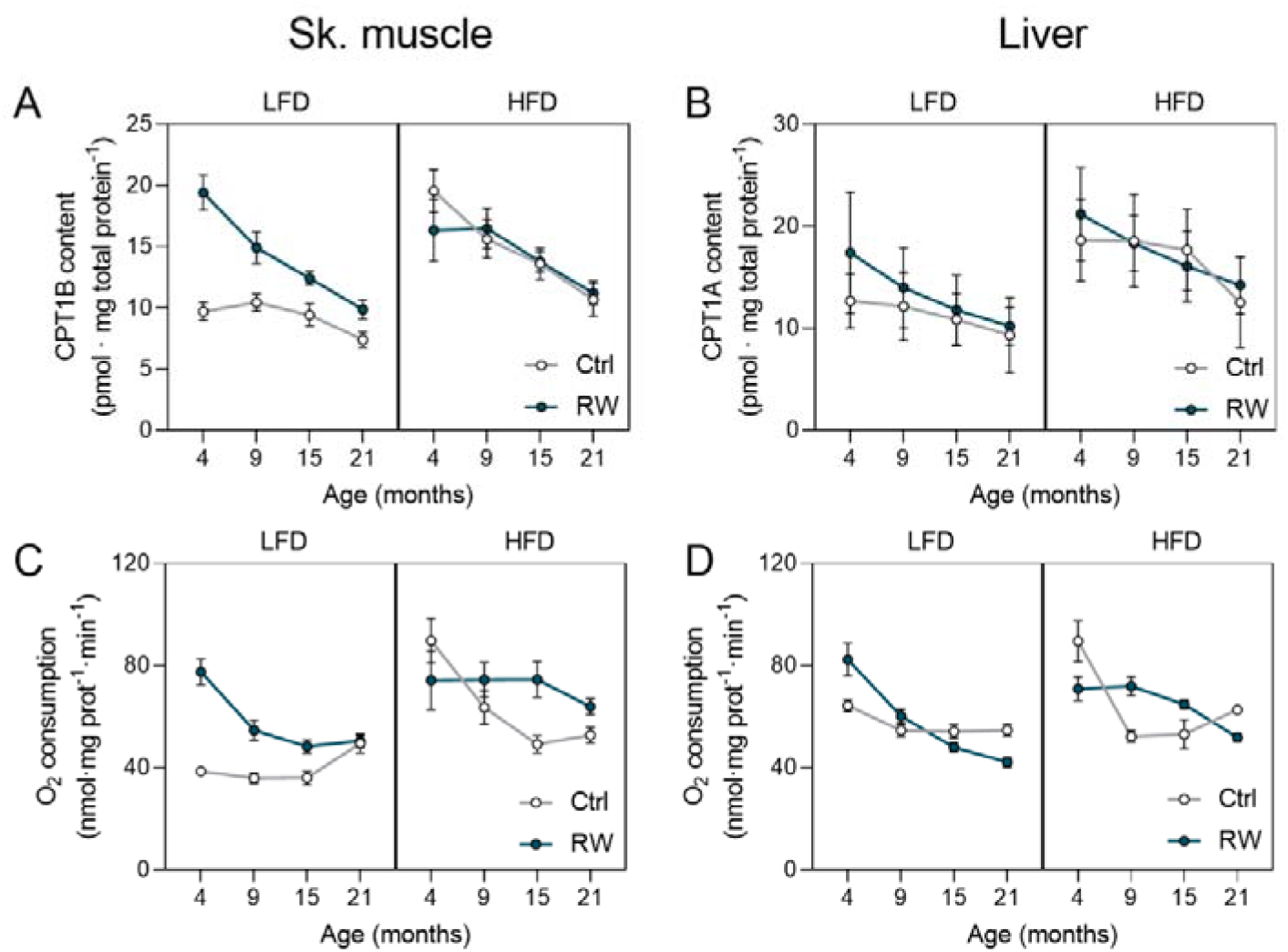
CPT1 content and oxidative capacity decline during aging. CPT1B protein content expressed per total skeletal muscle protein (A). CPT1A protein content expressed per total liver protein (B). Oxidative capacity (maximal ADP-stimulated O2 consumption) with the use of palmitoyl-CoA, carnitine and malate as substrates for skeletal muscle (C) and liver (D) expressed per total tissue protein. CPT1B protein content was available from Stolle et al., 2018. Skeletal muscle oxidative capacity data were reproduced from the same study (26). LFD: low-fat diet, HFD: high-fat high-sucrose diet, Ctrl: sedentary mice, RW: mice with voluntary access to a running wheel. Data are shown as mean ± SEM, n=4-8 per group.

In the liver, CPT1A (liver isoform) content was also increased by the HFD (p_diet_ < 0.05) and it exhibited a tendency of decline with age (Fig. 7B), even though there was no effect of the RW. A correlation between CPT1A and oxidative capacity was also observed in the liver, albeit weaker than in the muscle (Fig. 7D, S7B). In the liver, the oxidative capacity was decreased by age and enhanced by the HFD (p_age_, p_diet_ < 0.001), with no clear RW effect (Fig. 7D).

The steep decline of the IS-P between 4 and 9 months in the LFD groups (Fig. 6C) preceded the decline of CPT1B content of the muscle. Nevertheless, RW access increased CPT1B content and oxidative capacity predominantly in the 4 months old LFD group, the same group in which it rescued the IS-P the most. The correlation between CPT1B content and IS-P in the LFD group had an R^2^ of 0.55 (Fig. S7E). which indicates that other factors contributed 45% to the modulation of peripheral insulin sensitivity by age and RW. Altogether, the upregulation in β-oxidation capacity in the skeletal muscle by RW access only improved insulin sensitivity when there was no lipid overload (LFD groups).

## Discussion

Here, we dissected in detail how age, diet, physical activity, and their interaction affect peripheral and hepatic insulin sensitivity, and how these are associated with the mitochondrial fatty-acid oxidation capacity. To our knowledge, such a comprehensive analysis has never been conducted before. The most surprising result was that physical activity rescued the hepatic insulin sensitivity in aging mice, with only a minor age-dependent effect on the muscle. In addition, these beneficial effects of the RW were only observable in the LFD. In agreement with earlier results in the muscle (13), the HFD-dependent induction of CPT1 and mitochondrial fatty-acid oxidation was lost in aged mice, both in the muscle and in the liver. This loss of mitochondrial flexibility was not directly reflected in insulin sensitivity, suggesting that the latter also depends on other factors. Finally, RW access, which results in a chronic exercise regimen, rescued this age-dependent loss of CPT1B and oxidative capacity only in the skeletal muscle of LFD animals, the same diet group in which some improvement of peripheral insulin sensitivity was observed. An overview of the main findings is shown in Table 1.

**Table 1:** Overview of main findings and strength of associations*

| Parameter | Age (Old vs young) | Diet (HFD vs LFD) | Exercise (RW vs Ctrl) |
| --- | --- | --- | --- |
| **k<sub>2</sub> (clearance rate)** | Diet-dependent effect | ↓ | - |
| **EGP** | ↓↓ | - | ↓<br>↓ (LFD group) |
| **IS-P** | ↓↓<br>↓↓ (LFD group) | ↓↓↓ | ↑ |
| **IS-L** | ↓ | ↓↓ | ↑<br>↑ (LFD group) |
| **Oxidative capacity quadriceps** | ↓↓ | ↑↑↑ | ↑ |
| **Oxidative capacity liver** | ↓↓↓↓ | ↑ | Age-dependent effect |

\*Arrows represent the direction of change (increase vs decrease). The number of arrows represents to what extent a result is explained by each variable. 1 arrow: 0-10% variance, 2 arrows: 10-20% variance, 3 arrows: 20-30% variance, 4 arrows: >30% variance. Groups between parenthesis indicate that the variance is further explained by interaction with the specified group.

The computational model to analyze the tracer OGTT was inspired by the Oral Minimal Model for humans (37). A difference between mouse and human studies is that less frequent and smaller blood volumes can be obtained from mice (38). Recently, a more complex computational model was fitted to mouse tracer OGTT data that had been obtained with a similar experimental setup. The latter model included insulin-dependent and -independent glucose elimination (36). Although biologically more realistic, it is plausible that this higher complexity came at the expense of parameter identifiability (50), but this was not explicitly addressed (36). A limitation of this study is that glucose bioavailability was considered to be constant, irrespective of age, diet and physical exercise, even though this did not impact the relative differences between the groups. Moreover, the model does not accurately capture the initial dynamics of glucose absorption, given that transit compartments were not included.

Despite these simplifications, the calculated steady-state fluxes were comparable to those obtained from independent steady-state infusion experiments. A final limitation concerns the use of only male mice due to the large study size with 3 independent variables, since sex is known to play a role in the response to diet, age and exercise (51; 52).

Previously, peripheral insulin sensitivity has been defined as the ability of insulin to enhance glucose effectiveness (31). Since the latter is expressed by the apparent elimination rate constant *k_2_*, we defined the peripheral insulin sensitivity as IS-P = k_2_/INS_OGTT_. This parameter has the same units as the previously described insulin action parameter (31). IS-P is reminiscent of the MISI (muscle insulin sensitivity index) (5; 30). The MISI is defined as the slope of the glucose decay during the clearance phase of the OGTT divided by the average insulin concentration during the same period. The slope depends on glucose absorption and elimination, which in turn depends on the glucose concentration. In contrast to the glucose decay slope used to calculate the MISI, *k_2_* is specific for the elimination process and corrected for the glucose concentration. Similarly, the IS-L was adapted from an existing index for hepatic insulin sensitivity (29), the difference being that the IS-L analysis relies on the average response during the OGTT, whereas the classical analysis was based on fasting data. To capture the dynamics of the hepatic and peripheral insulin sensitivity during the OGTT would require more data points and additional assumptions, or multiple tracer experiments (45).

High-fat diets have long been used to induce obesity and insulin resistance in animal models (53). In agreement with this, the HFD decreased both IS-P and IS-L (Fig. 6C-D). In previous studies aging exacerbated HFD-induced insulin resistance (11; 12). In this study, this was confirmed by the HOMA-IR and it could be specifically attributed to the hepatic insulin response, as quantified by the IS-L. The peripheral insulin response (IS-P) was, however, equally low in all HFD groups, in contrast to our earlier findings that HFD-induced peripheral insulin resistance was aggravated by age (13). The studies differ, however, in terms of (i) diet duration, timing and composition, and (ii) mouse strain.

Despite the just mentioned differences in experimental setup, the CPT1B content and the mitochondrial capacity to oxidize fatty acids in the skeletal muscle recapitulate our previous findings (Fig. S7C-D). In both studies, older mice showed lower flexibility to upregulate the fatty-acid oxidation capacity in the muscle when fed a HFD (13). A similar pattern was observed in the liver (Fig. 7D). This mechanism was proposed to underlie the high muscle lipid accumulation in HFD-fed aged animals, consequently aggravating insulin resistance. In the present study, however, the IS-P decline with age preceded the drop in CPT1B content and fatty-acid oxidation capacity (Fig. 7A, C). Consequently, additional insulin-resistance-inducing mechanisms likely underlie the observed phenotype.

In this study, the RW was present throughout life and was not removed during the OGTT timeframe. However, since the mice did not exercise during the day, when the OGTT was conducted (Fig. S1), we here address the effects of chronic, and not acute, exercise on insulin sensitivity. The peripheral insulin sensitivity was barely enhanced by exercise in aged groups, which corroborates the findings by Short *et al*. in humans (24). However, several human studies have pointed to resistance or high-intensity aerobic training as strategies to improve insulin sensitivity in the elderly (3; 54; 55). Moreover, training regimens that result in weight loss could be a more effective strategy (56). In the present cohort, weight was maintained (26). Different explanations might underlie the heterogeneity of outcomes. First, the type and duration of exercise differed among studies. Second, the quantification of IR ranged from simple whole-body HOMA-IR quantification to more complex tracer and clamp studies. Third, mice run less as they grow older (57; 58), which was confirmed for the mice used here, especially for the HFD groups, and described in earlier reports about the same cohort (26; 35). Nevertheless, voluntary running promoted beneficial health adaptations, including mitochondrial remodeling (25; 26). We observed, however, that the insulin-desensitizing effects of the HFD outweighed the insulin-sensitizing response to endurance exercise. The net effect of diet and exercise on the insulin signaling cascade dictates glucose uptake rates and consequently insulin resistance development/prevention.

Lifelong RW access decreased the EGP in the LFD group (Fig. 5B, C). This corroborates previous findings after a short-term RW intervention in mice (32). Moreover, RW access intervention remarkably sustained a prolonged high hepatic insulin sensitivity during aging, albeit only in the LFD group. This was in stark contrast with the muscle, in which the RW did not prevent the age-dependent decline of the IS-P. The increase of the IS-L by RW access was not associated with CPT1A content or oxidative capacity. Rather, it may be attributed to the abovementioned increased capacity of hepatic glucose uptake *per se*, irrespective of insulin. Alternatively, it may derive from the action of insulin-sensitizing myokines, which can communicate the effect of exercise from the muscle to the liver (59).

In summary, based on a simple two-compartment model, EGP and muscle- and liver-specific insulin sensitivity could be reliably estimated from a non-invasive tracer-based OGTT. Hepatic and peripheral insulin sensitivity were both similarly decreased by aging and HFD. In LFD-fed mice, exercise enhanced peripheral insulin sensitivity mostly in young animals, and completely prevented the age-dependent loss of hepatic insulin sensitivity. These results, together with the applied methodology, may contribute to future research on the role of mitochondria in insulin resistance.

## Acknowledgments

We would like to thank Maaike Oosterveer and Angela Tol from the Laboratory of Pediatrics of the University Medical Center Groningen for valuable discussions.

## Author Contributions

MA.V.L., S.K. and B.M.B. developed the computational model, advised by F.A. and T.H.v.D. A. C.R., J.C. and A.T. conducted animal experiments. M.A.V.L., S.K. and A.C.R. conducted data analysis. T.H.v.D. analyzed steady-state labeling data. J.C.W. conducted proteomics experiments. C.J.V. and R.H.J.B. contributed with the experiments on bioavailability. A.K.G., D.J.R., G.J.v.D. and B.M.B. conceptualized and supervised the project. M.A.V.L. and B.M.B. wrote the manuscript. All authors read and approved the final manuscript.

## Guarantor Data Access and Responsibility Statement

B. M.B. is the guarantor of this work and, as such, had full access to all the data in the study and take responsibility for the integrity of the data and the accuracy of the data analysis. The authors declare no conflict of interest.

## Prior presentation

Parts of this study were presented orally at the Bioinformatics & Systems Biology Conference 2021, online, 15–16 June 2021 and at the International Study Group for Systems Biology 2022, Innsbruck – Austria, 19–23 September 2022.

