## Appendix 1 for "Age and diet modulate the insulin-sensitizing effects of exercise: a tracer-based oral glucose tolerance test"

**Appendix 1: Computational Model development**

The model consisted of three processes:

1. The import of glucose from the gastrointestinal compartment into the blood with rate *v_1_*;
2. The elimination of glucose from the plasma into the tissues and its subsequent storage or metabolism with rate *v_2_*; and
3. The metabolism or storage of glucose before it enters into the plasma pool, indicated as loss, with rate *v_L_*.

Rates *v* were expressed in µmol·kg^-1^·min^-1^.

The rates of these processes were described by first-order kinetics, with the following rate equations for the tracer:

$$v_{1}\left( q_{1} \right)= k_{1}\cdot q_{1} (1)$$

$$v_{L}\left( q_{1} \right)= k_{L}\cdot q_{1} (2)$$

$$v_{2}\left( c_{2} \right)= k_{2}\cdot c_{2}\cdot Vol (3)$$

All rate constants *k* were expressed in min^-1^. The conversion factor *Vol* in mL·kg^-1^ is the apparent distribution volume of the plasma compartment and was introduced to convert the concentration *c_2_* (mM) into a pool size (µmol·kg^‑1^), such that the rate *v_2_* is also expressed in µmol·kg^-1^·min^-1^.

Analogous rate equations for the unlabeled glucose were defined by:

$$v_{1}\left( Q_{1} \right)= k_{1}\cdot Q_{1} (4)$$

$$v_{L}\left( Q_{1} \right)= k_{L}\cdot Q_{1} (5)$$

$$v_{2}\left( C_{2} \right)= k_{2}\cdot C_{2}\cdot Vol (6)$$

Since tracer and unlabeled glucose exhibit chemically identical behavior, the rate constants *k_1_*, *k_2_*, and *k_L_* were assumed to be identical for both and were derived from the tracer kinetics.

In addition, unlabeled glucose is produced mainly by the liver, but also by the kidney and intestine. This process was quantified by the time‑dependent Endogenous Glucose Production, EGP*. Taken together, this led to the following set of ordinary differential equations (ODE):

$$\frac{dq_{1}}{dt}=-\left( k_{1}+k_{L} \right)\cdot q_{1} (7)$$

$$\frac{dc_{2}}{dt}=\frac{k_{1}\cdot q_{1}}{Vol}-k_{2}\cdot c_{2} (8)$$

$$\frac{dQ_{1}}{dt}=-\left( k_{1}+k_{L} \right)\cdot Q_{1} (9)$$

$$\frac{dC_{2}}{dt}=\frac{k_{1}\cdot Q_{1}}{Vol}{-k}_{2}\cdot C_{2}+{EGP}^{*}\left( t \right) (10)$$

Time *t* was expressed in min and EGP* in mM·min^-1^. The asterisk in EGP* denotes that it differs from the more commonly used EGP in µmol·kg^-1^·min^-1^.

The analytical solution of ODEs 7 and 8 gives (derived in the Supp. Material S2):

$c_{2}(t)= C\cdot\left( e^{-k_{2}t}-e^{-k_{a}t} \right)$	(11)

in which:

$$k_{a}=k_{1}+k_{L} (12)$$

The tracer time-course data were fitted to equation 11, yielding *C*, *k_2_* and *k_a_.* Identifiability and constraints are addressed in the Supp. Material S3.

To estimate the apparent volume of distribution of the tracer (*Vol,* the plasma compartment in the model), the bioavailability *F* is required. *F* is defined as:

$$F=\frac{k_{1}}{k_{1}+k_{L}} (13)$$

It can be derived (Supp. Material S2) that:

$$Vol= \frac{q_{1}(0)}{C}\cdot\frac{k_{a}\cdot F}{k_{a}-k_{2}}=\frac{q_{1}(0)}{C}\cdot\frac{k_{1}}{k_{a}-k_{2}} (14)$$

in which *q_1_*(0) represents the amount of tracer administered (1.6·10^3^ µmol·kg^-1^ in the present study).

The basal rate of disappearance for glucose *(*$R_{d}^{OGTT}$*)* in µmol·kg^-1^·min^-1^ was calculated according to the equation below:

$$R_{d}^{OGTT}= k_{2}\cdot C_{2}\left( 0 \right)\cdot Vol (15)$$

**EGP calculation**

The endogenous glucose production EGP* in mM·min^-1^ was computed by rearranging equation 10 as follows:

$${EGP}^{*}(t)=-\frac{k_{1}\cdot Q_{1}\left( t \right)}{Vol}+\frac{{dC}_{2}\left( t \right)}{dt}+k_{2}\cdot C_{2}\left( t \right) (16)$$

The components of the EGP can be determined as follows. From equation 9 it is derived that:

$$Q_{1}\left( t \right)= Q_{1}\left( 0 \right)\cdot e^{-\left( k_{1}+k_{L} \right)t}=Q_{1}\left( 0 \right)\cdot e^{-k_{a}t} (17)$$

in which *Q_1_(0)* is the amount of unlabeled glucose administered (3.9·10^3^ µmol·kg^-1^ in the present study). The compartment model presented here does not include transit compartments, since the absorption phase was too fast to capture more detail with our experimental setup.

Although *k_1_* and *Vol* in equation 16 depend on the choice of *F* (eq. 13-14), the ratio *k_1_/Vol* does not and by rearranging equation 14, it can be written as:

$$\frac{k_{1}}{Vol}= \frac{\left( k_{a}-k_{2} \right)\cdot C}{q_{1}\left( 0 \right)} (18)$$

Finally, *C_2_(t)* in equation 16 is obtained by fitting the time course of the unlabeled glucose concentration to:

$$C_{2}\left( t \right)=C_{0u}+C_{1u}\cdot e^{{-k}_{1u}t}+C_{2u}\cdot e^{-k_{2u}t} (19)$$

in which *u* denotes ‘unlabeled’ and the parameters *C_0u_*, *C_1u_*, *k_1u_*, *C_2u_*, and *k_2u_* are considered phenomenological constants without a direct relation to the underlying rate equations. Therefore, unique identifiability of the time parameters is of less relevance than a good fit of the data. Nevertheless, to aid the identification of biologically pertinent fits, *k_1u_* was fixed at the same value obtained for *k_a_* per animal based on tracer fits. Finally, taking the time derivative of equation 19 gives:

$$\frac{{dC}_{2}\left( t \right)}{dt}={-C}_{1u}\cdot k_{1u}\cdot e^{{-k}_{1u}t}-C_{2u}\cdot{k_{2u}\cdot e}^{{-k}_{2u}t} (20)$$

To compute the EGP* in mM·min^-1^, equation 16 can be rewritten in full, which is independent of the bioavailability *F*:

$${EGP}^{*}(t)=C_{1u}\cdot\left( k_{2}-k_{1u} \right)\cdot e^{{-k}_{1u}\cdot t}-\frac{\left( k_{a}-k_{2} \right)\cdot C}{q_{1}\left( 0 \right)}\cdot Q_{1}\left( 0 \right)\cdot e^{{-k}_{a}\cdot t}$$

$$+C_{2u}\cdot\left( k_{2}-k_{2u} \right)\cdot e^{{-k}_{2u}\cdot t} + k_{2}\cdot C_{0u} (21)$$

To convert EGP* (mM·min^-1^) to EGP (µmol·kg^-1^·min^-1^), EGP* was multiplied by *Vol*, which is a function of the bioavailability *F* (equation 14).

**Steady-state intravenous infusion (SS‑IV)**

Experiments were performed according to the protocol described by van Dijk et al. (1). Animals were fasted overnight for 9h. On the day of the experiment, animals were placed in cages without access to the RW. Animals were continuously infused with [U-^13^C_6_]-glucose for 6 hours, and blood glucose was sampled every hour. Glucose and tracer measurements were conducted as described above. Calculation of $R_{d}^{SSIV}$ and ${EGP}^{SSIV}$was conducted as described below:

$$\frac{dc_{2}(SSIV)}{dt}=\frac{v_{infusion}-v_{consumption}}{Vol} (22)$$

$$v_{infusion}=R_{inf}\cdot M_{I} (23)$$

$$v_{consumption}=R_{d}^{SSIV}\cdot M_{C} (24)$$

Where *R_inf_* represents the glucose infusion rate in µmol·kg^-1^·min^-1^, *M_I_* is the fractional abundance of tracer in the infusion solution, *M_C_* is the fractional abundance of tracer measured in the circulation after correction for natural abundance, and $R_{d}^{SSIV}$ is the steady-state rate of disappearance of total glucose in µmol·kg^-1^·min^-1^. At steady state:

$$\frac{dc_{2}\left( SSIV \right)}{dt}=0 (25)$$

$$\therefore R_{d}^{SSIV}=\frac{R_{inf}\cdot M_{I}}{M_{C}} (26)$$

Following the same logic, the rates of appearance *(EGP^SS‑IV^ + R_inf_)* and disappearance *(*$R_{d}^{SSIV}$*)* for total glucose (unlabeled + tracer) should be equal to each other at steady state, resulting in:

$${EGP}^{SSIV}=R_{d}^{SSIV}-R_{inf} (27)$$

in which all rates, including *EGP^SS‑IV^*, are in µmol·kg^-1^·min^-1^.

**Insulin sensitivity indices**

HOMA-IR was calculated according to (2):

$$HOMAIR= \frac{C_{2}\left( 0 \right)\cdot INS(0)}{14.1} (28)$$

in which *C_2_(0)* represents fasting glucose levels in mM, *INS(0)* fasting insulin levels in mU·L^‑1^, and 14.1 the reference value for mice in mM· mU·L^-1^.

Based on the OGTT data, the insulin sensitivity index for peripheral tissues (IS-P) was defined as:

$$ISP=\frac{k_{2}}{{INS}_{OGTT}} (29)$$

in which *INS_OGTT_* is the average insulin level over all time points during the OGTT time frame, calculated for each animal separately.

The adapted liver insulin sensitivity (IS-L) was defined as:

$$ISL=\frac{\bar{EGP}\times\bar{INS}_{OGTT}}{EGP\times{INS}_{OGTT}} (30)$$

in which *EGP* is the time-averaged EGP from 5 to 120 min. The denominator represents a value calculated per specific group, whereas the numerator is the average *EGP* times average *INS_OGTT_* (over all animals in all 16 experimental groups). With this, the mean was centered and a dimensionless index was obtained.

All indices were obtained by using the mean *C_2_(0)*, *k_2_*, *EGP*, *INS(0)* and *INS_OGTT_* values per experimental group. To obtain the distribution (SD and SEM) of the calculated indices, the distribution of the product or ratio of the means was calculated assuming normal distribution of the data.
