## Supplemental Materials and Figures for "Age and diet modulate the insulin-sensitizing effects of exercise: a tracer-based oral glucose tolerance test"

**Supplemental Material 1: Natural abundance correction**

***Definitions***

*m_0_*	The nominal mass of the compound ion, rounded to the nearest integer, with only the most abundant isotopes (e.g.${}_{1}^{1}H$, ${}_{6}^{12}C$, ${}_{7}^{14}N$)

*m_i_*	The mass of the compound ion incremented by *i* above the nominal mass, due to incorporation of *i* isotopes with a higher mass than the most abundant (*e.g.*${}_{1}^{2}H$, ${}_{6}^{13}C$, ${}_{7}^{15}N$)

*A_i_*	The recorded absolute intensity (peak area) in the mass spectrum of a compound ion at $m/z$=*m_i_*

*Isotopologues*	Molecular species that have the same chemical formula, but differ in isotopic content and therefore in mass

*Tracer*	Here: stable-isotope labeled tracer, a compound in which one or more atoms have been replaced by a stable isotope, which allows the compound and its biochemical products to be distinguished from endogenous metabolites

ID	Isotopologue distribution, consisting of the fractional contributions of the masses of interest of the particular compound ion

***K***	The representation of the *measured* ID as a vector

***M***	The representation of the ID as a vector, *after correction* for natural abundance of isotopes

*K_i_*	Element of the vector ***K*** representing the *measured* fractional abundance of the isotopologue with mass *m_i_*

*M_i_*	Element of the vector ***M*** representing the fractional abundance of the isotopologue with mass *m­_i_* *after correction* of *K_i_* for the contribution of naturally occurring isotopes

***L***	The correction matrix that converts the measured fractional abundance vector ***K*** into the corrected matrix ***M***

$L_{j}^{i}$	Element $L_{j}^{i}$ of the correction matrix ***L***, denoting the probability that an isotopologue in which *i* atoms were substituted for their corresponding stable isotope from the tracer, is heavier by an additional *j* mass units due to incorporation of naturally occurring isotopes.

***General procedure***

An isotopologue of mass *m_i_* has a mass increment *i* above the nominal mass of the compound. The mass increment can originate from the tracer or from naturally occurring isotopes. A correction is necessary to transform the measured isotopologue distribution (ID) into the distribution that should have been observed had there been no naturally occurring isotopes. The method described here is based on the *measured* ID of the unlabeled, baseline sample that was obtained prior to addition of the stable isotope labeled tracer. What is new, at least to our knowledge, is the use of this *measured* ID of the baseline sample to reconstruct the skewed correction matrix ***L*** (1; 2). We developed this method for compounds with a low fractional enrichment (typically less than 5%) in combination with a high molecular mass, such as metabolites that have been derivatized for GC-MS measurements. In these cases the generally accepted deviation of the enrichment of the baseline sample after correction (± 0.4%), based on the default error for GC-MS measurements (3), would be substantial relative to the enrichment due to label incorporation. The general part of the procedure works as follows.

The vector ***K*** is the measured ID, defined as:

$$\boldsymbol{K}=\left[ \begin{matrix} \begin{matrix} K_{0} \\ K_{1} \\ K_{2} \end{matrix} \\ \ldots\\ K_{n} \end{matrix} \right]$$

in which *n* is the number of atoms that can be substituted with a heavy isotope, and:

$$K_{i}= {A_{i}}/{\sum_{i=0}^{n} A_{i}}$$

Analogously a vector ***M*** of corrected fractional abundances is defined as:

$$\boldsymbol{M}=\left[ \begin{matrix} \begin{matrix} M_{0} \\ M_{1} \\ M_{2} \end{matrix} \\ \ldots\\ M_{n} \end{matrix} \right]$$

in which element *M_i_* represents the fractional abundance of the isotopologue with mass *m_i_* after correction for natural isotope abundance, i.e. the fractional abundance that can be attributed to the tracer. ***M*** is calculated from ***K*** according to:

$$\boldsymbol{K}=\boldsymbol{L\cdot M}$$

in which

$$\boldsymbol{L}=\left[ \begin{matrix} L_{0}^{0} & 0 & 0 & \ldots& 0 \\ L_{1}^{0} & L_{0}^{1} & 0 & \ldots& 0 \\ L_{2}^{0} & L_{1}^{1} & L_{0}^{2} & \ldots& 0 \\ \ldots& \ldots& \ldots& \ldots& 0 \\ L_{n}^{0} & L_{n-1}^{1} & L_{n-2}^{2} & \ldots& L_{0}^{n} \end{matrix} \right]$$

The elements $L_{j}^{i}$ of the correction matrix ***L*** denote the probability that an isotopologue in which *i* atoms were incorporated from the stable-isotope labeled substrate, is heavier by an additional *j* mass units due incorporation of naturally occurring isotopes.

The elements of the first column of ***L*** *(*$L_{j}^{0}$**)** denote the probabilities that measured isotopologues are exclusively labeled by naturally occurring isotopes, as would be the case in the baseline sample prior to addition of the tracer. When one atom has been replaced by a heavy isotope from the tracer, the number of atoms of the compound that can be substituted with a naturally occurring isotope is *n-1*. If *i* atoms are substituted by isotopes originating from the labeled substrate, then the other *n‑i* atoms have a probability of being labeled by naturally occurring isotopes. Since this probability depends on the number of the other *n-i* unlabeled atoms, the ***L*** matrix becomes ‘skewed’ (1; 2), i.e. the columns are not just shifted relative to each other, but the probabilities differ, *i.e.* $L_{j}^{i}\neq L_{j}^{i+1}$.

If matrix ***L*** is known, vector ***M*** is calculated as follows.

Given: ***K*** = ***L***·***M***. Then: ***L^-1^***·***K* = *L^-1^***·***L***·***M*** and thus: ***L^-1^***·***K* = *M****.* Hence, to solve ***M***, matrix ***L*** is inverted and the resulting matrix ***L^-1^*** multiplied to vector ***K***.

Most often, matrix ***L*** is calculated from the known abundances of naturally occurring isotopes (4). In cases with a high enrichment with isotopes from the tracer, this approach is sufficiently accurate. It may become problematic for large molecules with a low isotope enrichment from the tracer. In these cases the generally accepted deviation of the enrichment of the baseline sample after correction (± 0.4%) would be substantial relative to the enrichment due to label incorporation. Then, the baseline sample provides an independent measurement of the actual ID due to naturally occurring isotopes. There are, however, no measured IDs of the compound with *i* tracer isotopes atoms incorporated, due to a lack of such standards. This implies that there is only information for the first column of ***L***. Below, we derive a method to construct the skewed ***L*** matrix, based on the measured baseline sample.

***Constructing the L matrix from the measured baseline samples***

To compute the skewed ***L*** matrix from the measured ID of the baseline sample, we were inspired by (5), who derived the ID (vector ***K*)** for a molecule composed of two chemical fragments *X* and *Y* with mutually independent isotopologue distributions. The isotopologue distribution (*K_0_, K_1_, K_2_,* …) of a molecule consisting of a fragments *X* with isotopologue distribution (*p_0_*, *p_1_*, *p_2_*, …) and a fragment *Y* with isotopologue distribution (*q_0_, q_1_, q_2_*, …) can be computed according to:

$$\left[ \begin{matrix} p_{0} & 0 & 0 & \ldots& \ldots\\ p_{1} & p_{0} & 0 & \ldots& \ldots\\ p_{2} & p_{1} & p_{0} & \ldots& \ldots\\ p_{3} & p_{2} & p_{1} & \ldots& \ldots\\ \ldots& \ldots& \ldots& \ldots& \ldots\\ p_{n} & p_{n-1} & p_{n-2} & \ldots& \ldots\end{matrix} \right]\cdot\left[ \begin{matrix} q_{0} \\ q_{1} \\ q_{2} \\ \ldots\\ \ldots\end{matrix} \right]=\left[ \begin{matrix} K_{0} \\ K_{1} \\ K_{2} \\ \ldots\\ \ldots\end{matrix} \right]$$

Note that here we do not have a skewed matrix: the columns containing *p_0_ – p_n_* are just shifted relative to each other, as the equation describes different combinations of the same fragments. We used the same principle here. The measured isopologue was considered to consist of two parts i.e. *T^i*^* and *R^-i^*. Part *T^i*^*, the ‘tracer part of the compound’, accounts for the incorporated isotopes from the tracer, while part *R^-i^*, the ‘rest of the molecule’, accounts for the incorporation of naturally occurring isotopes of all atoms in the rest of the molecule. This tracer part *T^i*^* is not a chemical fragment of the molecule, but just the total of the heavy elements from the tracer. The asterisk denotes that this is the labeled version of *T^i*^*, since we will later distinguish it from the equivalent part of the molecule *T^i^* without the tracer isotopes incorporated. The natural isotopologue distribution vector ***R^-i^*** of the fragment *R^-i^* can be written as:

$$\boldsymbol{R}^{\boldsymbol{-i}}=\left[ \begin{matrix} R_{0}^{-i} \\ R_{1}^{-i} \\ R_{2}^{-i} \\ \ldots\\ R_{n-i}^{-i} \end{matrix} \right]$$

in which element $R_{j}^{-i}$ denotes the probability that part *R^-i^* lacking *i* labeled atoms, has a mass increment of *j* due to incorporation of naturally occurring isotopes. The correction matrix ***L*** can now be constructed according to:

$$\boldsymbol{L}=\left[ \begin{matrix} R_{0}^{0} & 0 & 0 & \ldots& 0 \\ R_{1}^{0} & R_{0}^{-1} & 0 & \ldots& 0 \\ R_{2}^{0} & R_{1}^{-1} & R_{0}^{-2} & \ldots& 0 \\ \ldots& \ldots& \ldots& \ldots& 0 \\ R_{n}^{0} & R_{n-1}^{-1} & R_{n-2}^{-2} & \ldots& R_{0}^{-n} \end{matrix} \right]$$

The first column ***R^0^*** (*i* = 0) is the isotopologue distribution of the unlabeled baseline sample prior to addition of the tracer (vector ***K^0^***). The key question is how to identify the isotopologue distributions ***R^-i^*** for *i ≠ 0.*

To obtain ***R^-i^*** we have to start with the ID of the baseline sample. The ID of the unlabeled compound in a part *T^i^* (the unlabeled equivalent of *T^i*^*) and a part *R^-i^*. Suppose that we have a compound X*_k_*Y*_l_*Z*_m_* with *n* atoms X that can be substituted for a stable isotope from the tracer. The compound can be perceived to consist of a part *T^i^*, equivalent to *X_i_*, and a remainder *R^-i^*, equivalent to X*_k-i_*Y*_l_*Z*_m_*, with *i ∈* [1 – *n*]. The probability $s_{j}^{i}$ that *T^i^* contains *j* heavy atoms due to natural abundance, is calculated according to:

$$s_{j}^{i}=\frac{i!}{j!\cdot\left( i-j \right)!}\cdot\left( t_{0} \right)^{i-j}\cdot\left( t_{1} \right)^{j}$$

in which *t_1_* is the known fractional natural abundance of the stable isotope of atom *X* and *t_0_* = 1 ‑ *t_1_*. Since the isotopologue distributions of *T^i^* and *R^-i^* are mutually independent, according to Lee (5) we can write:

$$\underset{\boldsymbol{T}^{\boldsymbol{i}}}{\underbrace{\left[ \begin{matrix} s_{0}^{i} & 0 & 0 & \ldots& 0\ldots\\ s_{1}^{i} & s_{0}^{i} & 0 & \ldots& 0\ldots\\ s_{2}^{i} & s_{1}^{i} & s_{0}^{i} & \ldots& 0\ldots\\ \ldots& \ldots& \ldots& \ldots& 0\ldots\\ s_{i}^{i} & s_{i-1}^{i} & s_{i-2}^{i} & \ldots& s_{0}^{i} \end{matrix} \right]}}\cdot\underset{\boldsymbol{R}^{\boldsymbol{-i}}}{\underbrace{\left[ \begin{matrix} R_{0}^{-i} \\ R_{1}^{-i} \\ R_{2}^{-i} \\ \ldots\\ R_{n-i}^{-i} \end{matrix} \right]}=}\underset{\boldsymbol{K}^{\boldsymbol{0}}}{\underbrace{\left[ \begin{matrix} K_{0}^{0} \\ K_{1}^{0} \\ K_{2}^{0} \\ \ldots\\ K_{n}^{0} \end{matrix} \right]}}$$

in which vector ***K^0^*** denotes the measured ID of the baseline sample. Thus, for the unlabeled sample we can dissect the measured isotopologue vector ***K^0^*** into:

$$\boldsymbol{T}^{\boldsymbol{i}}\boldsymbol{\cdot}\boldsymbol{R}^{\boldsymbol{-i}}=\boldsymbol{K}^{\boldsymbol{0}}$$

Therefore: $\left[ \boldsymbol{T}^{\boldsymbol{i}} \right]^{\boldsymbol{-1}}\boldsymbol{\cdot T}^{\boldsymbol{i}}\boldsymbol{\cdot}\boldsymbol{R}^{\boldsymbol{-i}}=\left[ \boldsymbol{T}^{\boldsymbol{i}} \right]^{\boldsymbol{-1}}\boldsymbol{\cdot}\boldsymbol{K}^{\boldsymbol{0}}$ and thus: $\boldsymbol{R}^{\boldsymbol{-i}}=\left[ \boldsymbol{T}^{\boldsymbol{i}} \right]^{\boldsymbol{-1}}\boldsymbol{\cdot}\boldsymbol{K}^{\boldsymbol{0}}$*.* Hence, to solve $\boldsymbol{R}^{\boldsymbol{-i}}$, matrix $\boldsymbol{T}^{\boldsymbol{i}}$ is inverted and multiplied to vector $\boldsymbol{K}^{\boldsymbol{0}}$.

This calculation is repeated *n* times, each time with a new matrix $\boldsymbol{T}^{\boldsymbol{i}}$ to calculate $\boldsymbol{R}^{\boldsymbol{-i}}$ for each *i*. In this way *n* vectors $\boldsymbol{R}^{\boldsymbol{-i}}$ with elements $R_{j}^{-i}$ are calculated. The vectors ***K^0^*** and ***R^-i^*** are then inserted as columns to form the ***L*** matrix with$L_{j}^{o}$ = $K_{j}^{0}$ and $L_{j}^{i}$ = $R_{j}^{-i}$.

In the current study a deuterium label has been used and the intensity of isotopologues with masses *m_0_ – m_4_* have been measured. Therefore, *X* = ${}_{1}^{1}H$ and *n* = 4, so *i* $\in$ [0 – 4]. The fractional natural abundance of ${}_{1}^{2}H$ has been reported to be 0.0115 % (6), thus *t_1_* = 0.000115. In the accompanying Excel sheet (Supplemental File 1) the calculations are done for [6,6-^2^H_2_]-glucose measured by GC-MS as the penta-acetate derivative.

Finally, why is this method more appropriate for samples with a very low label enrichment in large molecules? If we derive the ***L*** matrix from natural isotope abundances (the state-of-the art method), for a large molecule small errors in individual isotope abundances may accumulate to a relatively large error. In the here described method, the natural abundances are used only to compute the ID of the small part *T^i^* and thus the cumulative error is much smaller. We checked that in our dataset the state-of-the-art method gave rise to deviations of less than 0.4% in the baseline sample, indicating that the quality of the measurements was good.

**Supplemental Material 2: Analytical solutions of simple tracer model**

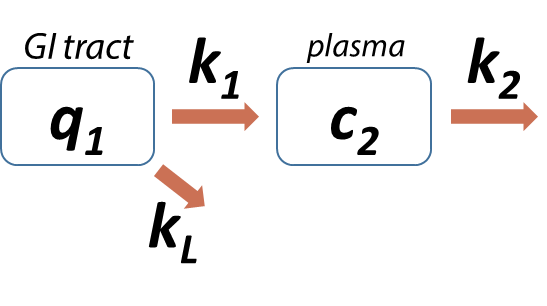

**Figure A1:** Two-compartment model of tracer kinetics

To derive general characteristics of the tracer kinetics, the compartment model depicted in Figure A1 was solved analytically. *q_1_* and *c_2_* represent the amount and concentration of the tracer in compartment *1* and *2* respectively. In the case of an oral gavage, *q_1_* represents a gastrointestinal compartment (µmol·kg^-1‑^) and *c_2_* the plasma compartment (mM). Rate constant *k_L_* refers to the loss of tracer before it reaches the plasma compartment.

The system is described by the following set of ordinary differential equations (ODE, cf. Research Design and Methods section for a motivation of the model equations):

$\frac{dq_{1}}{dt}=-\left( k_{1}+k_{L} \right)\cdot q_{1}$	(Eq. S2.1)

$\frac{dc_{2}}{dt}=\frac{k_{1}\cdot q_{1}}{Vol}{-k}_{2}\cdot c_{2}$	(Eq. S2.2)

In matrix notation, this is equivalent to:

$\frac{d}{dt}\left[ \begin{matrix} q_{1} \\ c_{2} \end{matrix} \right]=\left[ \begin{matrix} -\left( k_{1}+k_{L} \right) & 0 \\ +k_{1}/Vol & -k_{2} \end{matrix} \right]\cdot\left[ \begin{matrix} q_{1} \\ c_{2} \end{matrix} \right]$	(Eq. S2.3)

which can be written as:

$\dot{\boldsymbol{x}}=\boldsymbol{A}\cdot\boldsymbol{x}$	(Eq. S2.4)

This has the general solution:

$\boldsymbol{x}=s_{1}\cdot e^{\lambda_{1}t}\cdot\boldsymbol{u}_{1}{+ s}_{2}\cdot e^{\lambda_{2}t}\cdot\boldsymbol{u}_{2}$	(Eq. S2.5)

in which *λ_1_* and *λ_2_* are the eigenvalues of matrix ***A,*** and ***u_1_*** and ***u_2_*** the corresponding eigenvectors.

The eigenvalues of ***A*** are calculated from:

$det\left( \boldsymbol{A}-\lambda\boldsymbol{I} \right)=0$	(Eq. S2.6)

and:	$\left( \boldsymbol{A}-\lambda\boldsymbol{I} \right)\cdot\boldsymbol{u}=\boldsymbol{0}$	(Eq. S2.7)

This yields eigenvalues *λ _1_* = -(*k_1_+k_L_)* and *λ _2_* = -*k_2_* with eigenvectors:

$\boldsymbol{u}_{\boldsymbol{1}}=\left[ \frac{\begin{matrix} 1 \\ k_{1} \end{matrix}}{Vol\cdot\left( k_{2}-k_{1}-k_{L} \right)} \right]$ and $\boldsymbol{u}_{\boldsymbol{2}}=\left[ \begin{matrix} 0 \\ 1 \end{matrix} \right]$	(Eq. S2.8)

This leads to:	$q_{1}= s_{1}\cdot e^{-\left( k_{1}+k_{L} \right)t}$	(Eq. S2.9)

$c_{2}= \frac{s_{1}}{Vol}\cdot\frac{k_{1}}{k_{2}-k_{1}-k_{L}}\cdot e^{-\left( k_{1}+k_{L} \right)t}+s_{2}\cdot e^{-k_{2}t}$	(Eq. S2.10)

With:	*q_1_*(0) *= q_1,0_* and *c_2_*(0) = 0, the solution becomes:

$q_{1}= q_{1,0}\cdot e^{-\left( k_{1}+k_{L} \right)t}$	(Eq. S2.11)

$c_{2}= \frac{q_{1,0}}{Vol}\cdot\frac{k_{1}}{k_{2}-k_{1}-k_{L}}\cdot e^{-\left( k_{1}+k_{L} \right)t}-\frac{q_{1,0}}{Vol}\cdot\frac{k_{1}}{k_{2}-k_{1}-k_{L}}\cdot e^{-k_{2}t}$	(Eq. S2.12)

In pharmacokinetics, the bioavailability of the tracer, *i.e.* the fraction that reaches the plasma compartment, is computed as the ratio between the area under the *c_2_* curve for an oral versus an intravenous administration.

For the oral administration as solved above, the area under the curve for *c_2_* becomes:

$${AUC}_{oral}=\int_{0}^{\infty} c_{2}\left( t \right)dt=\frac{q_{1,0}}{Vol}\cdot\frac{k_{1}}{k_{2}-k_{1}-k_{L}} \int_{0}^{\infty} \left( e^{-\left( k_{1}+k_{L} \right)t}-e^{-k_{2}t} \right) dt$$

$=\left( \frac{q_{1,0}}{Vol\cdot k_{2}} \right)\cdot\frac{k_{1}}{k_{1}+k_{L}}$	(Eq. S2.13)

Had the same amount of tracer been administered intravenously (IV), *i.e*. directly into the plasma compartment, the kinetics would be described by:

$\frac{dc_{2}}{dt}={-k}_{2}\cdot c_{2}$	(Eq. S2.14)

with:	$c_{2}\left( 0 \right)=\frac{q_{1,0}}{Vol}$	(Eq. S2.15)

which leads to:

$c_{2}= \frac{q_{1,0}}{Vol}\cdot e^{-k_{2}t}$	(Eq. S2.16)

Now the area under the curve becomes:

${AUC}_{IV}=\int_{0}^{\infty} c_{2}(t)dt=\frac{q_{1,0}}{Vol} \int_{0}^{\infty} e^{-k_{2}t}dt=\frac{q_{1,0}}{Vol\cdot k_{2}}$	(Eq. S2.17)

And thus:

$F=\frac{{AUC}_{oral}}{{AUC}_{IV}}=\frac{k_{1}}{k_{1}+k_{L}}$	(Eq. S2.18)

This is a logical outcome, since it corresponds to the fraction of *q_1_* that is transported to compartment 2.

Based on the analytical solution above, equation S2.12 describing the tracer kinetics can be simplified to:

$c_{2}(t)= C\cdot\left( e^{-k_{2}t}-e^{-k_{a}t} \right)$	(Eq. S2.19)

in which:	$k_{a}=k_{1}+k_{L}$	(Eq. S2.20)

and thus:	$k_{1}=k_{a}\cdot F$	(Eq. S2.21)

$C= -\frac{q_{1}\left( 0 \right)}{Vol}\cdot\frac{k_{1}}{k_{2}-k_{1}-k_{L}}=-\frac{q_{1}\left( 0 \right)}{Vol}\cdot\frac{k_{a}\cdot F}{k_{2}-k_{a}}$	(Eq. S2.22)

Thus, if *C*, *k_a_* and *k_2_* are fitted from the data, the apparent distribution volume *Vol* can be calculated from the bioavailability *F* or *vice versa*. We are aware that we have derived here only classical pharmacokinetics equations, but it clarified (1) that the *k_a_* fitted to tracer kinetics is only an apparent absorption rate constant, while the actual rate constant *k_1_* equals *k_a_*·*F,* and (2) what is the basis for the calculation of the apparent volume *Vol* from *F*. The latter is used to express the endogenous glucose production (EGP) in μmol·min^-1^·kg^-1^.

**Supplemental Material 3: Data fitting and identifiability**

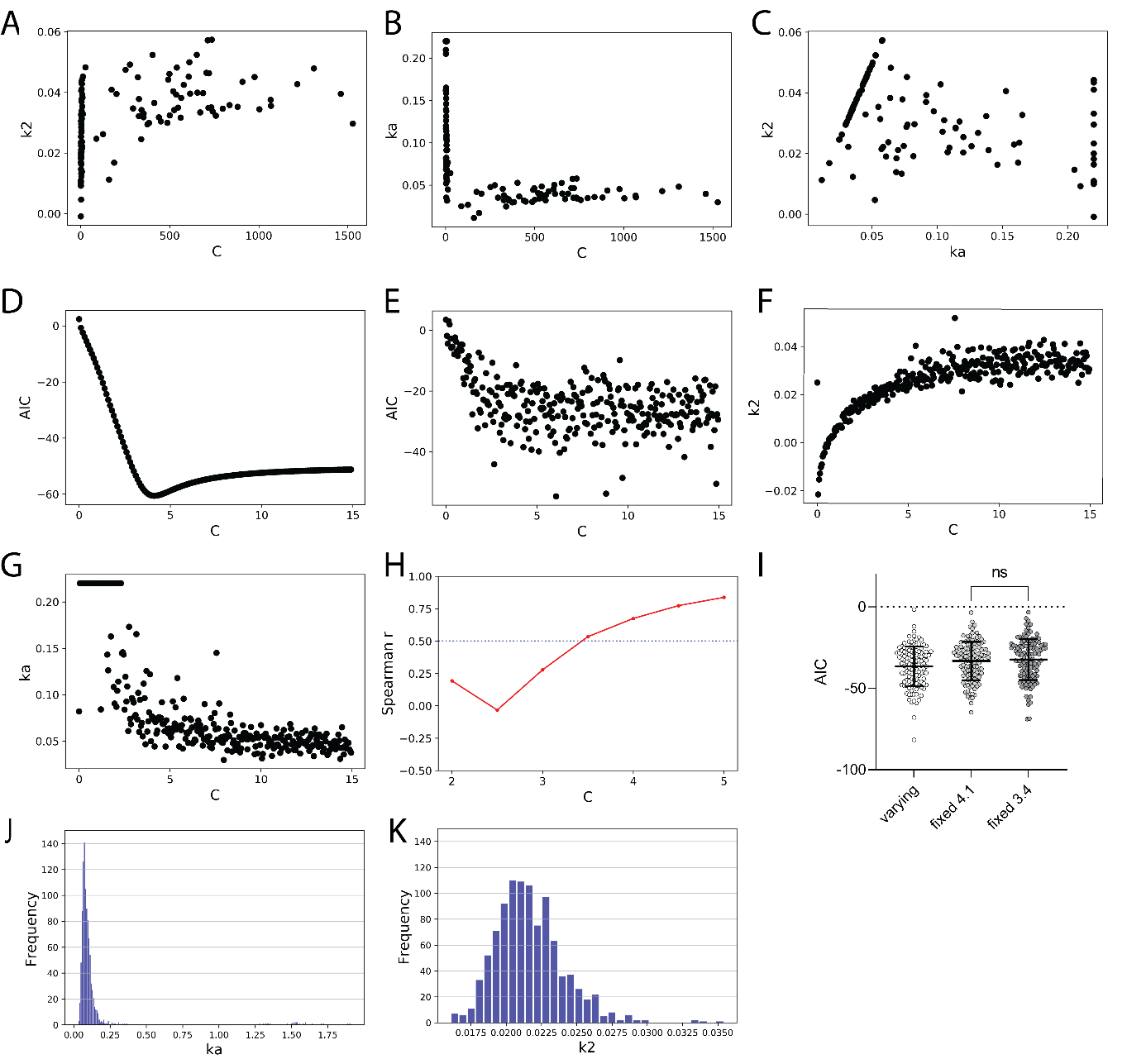

**Figure A2:** Correlations between all 3 parameters obtained from the tracer fit for all animals, with *k_a_* with an upper bound of 0.22 (A, B, C). AIC values (D) derived from fitting of data values for a fixed *C* ranging from 0 and 15 for the mean values per time point. AIC (E), *k_2_* (F) and *k_a_* (G) values derived in which a random variation was included in the mean values according to the distribution of the original dataset (H). Spearman correlation coefficient *r* between *k_a_* and *k_2_* for model fit with different fixed values for *C*. Data were randomly generated according to mean and distribution for all time points (H). AIC comparison for model fits with the use of data from all animals. *C* was let to vary or was fixed at 4.1 or 3.4 (I). Expected distribution for *k_a_* (J) and *k_2_* (K) from 1,000 fits based on the distribution of data randomly generated according to mean and distribution for all time points.

To obtain *C*, *k_a_* and *k*_2_, the time course of tracer data for each animal (mM) was fitted to equation 11 $\left[ q_{2}(t)= C\cdot\left( e^{-k_{2}t}-e^{-k_{a}t} \right) \right]$. Not all three parameters were identifiable from the tracer data due to collinearity (Fig. A2A-C). Given the intrinsic biological information contained in *k_a_* and *k_2_*, *C* was selected to be fixed at 3.4 for the fits, based on (i) the lowest AIC (Akaike’s Information Criterion for model selection) for data mean per time point (Fig. A2D, E), and on (ii) a Spearman correlation coefficient between *k_a_* and *k_2_* lower than 0.5, based on 1 000 simulations of synthetic data (Fig. A2H). These assumptions did not interfere with the obtained AIC for the data fit (Fig. A2I). The constant *k_a_* was constrained within the 95% CI of the median (0.047 – 0.22 min^-1^) to avoid biologically inconsistent outliers, based on the expected distribution from synthetic data (Fig. A2J, K). The constant *k_2_* was not constrained. Within the region of C values resulting in low AIC, *k_a_* and *k_2_* did not heavily depend on the choice of *C* (Fig. A2F, G).

The lack of correlation between *k_a_* and *k_2_* (Fig A3A) implies that both could be independently identified. The use of a more complex model including an extra distribution compartment and consequently a second elimination term did not result in better fits of the data and impaired identification of the elimination constant(s) (Fig. A3B, C). This is in agreement with analysis of human OGTT data, in which even more time points were taken due to the larger blood volume, and yet one elimination compartment was sufficient to fit the data accurately (7).

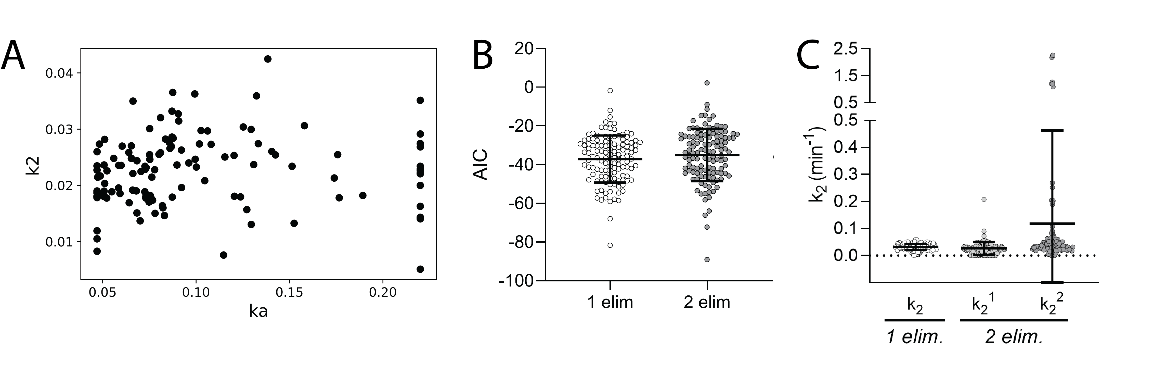

**Figure A3:** Scatter plot for *k_a_* and *k_2_* values for fits with fixed *C* at 3.4 and applied boundaries for *k_a_* (A). Comparison for all animals in the dataset when fitting tracer data using either 1 elimination term [$q_{2}(t)= C\cdot\left( e^{-k_{2}t}-e^{-k_{a}t} \right)]$ or 2 elimination terms ${[q}_{2}\left( t \right)= C_{1}\cdot e^{-k_{2}^{1}\cdot t}+C_{2}\cdot e^{-k_{2}^{2}\cdot t}-(C_{1}+C_{2})\cdot e^{-k_{a}t}]$ for AIC (B) and for elimination constants (C). A second elimination constant did not improve the AIC. Moreover, the second elimination constant could not be estimated, as is obvious from the large spread in the fitted values.

**Supplemental figures**

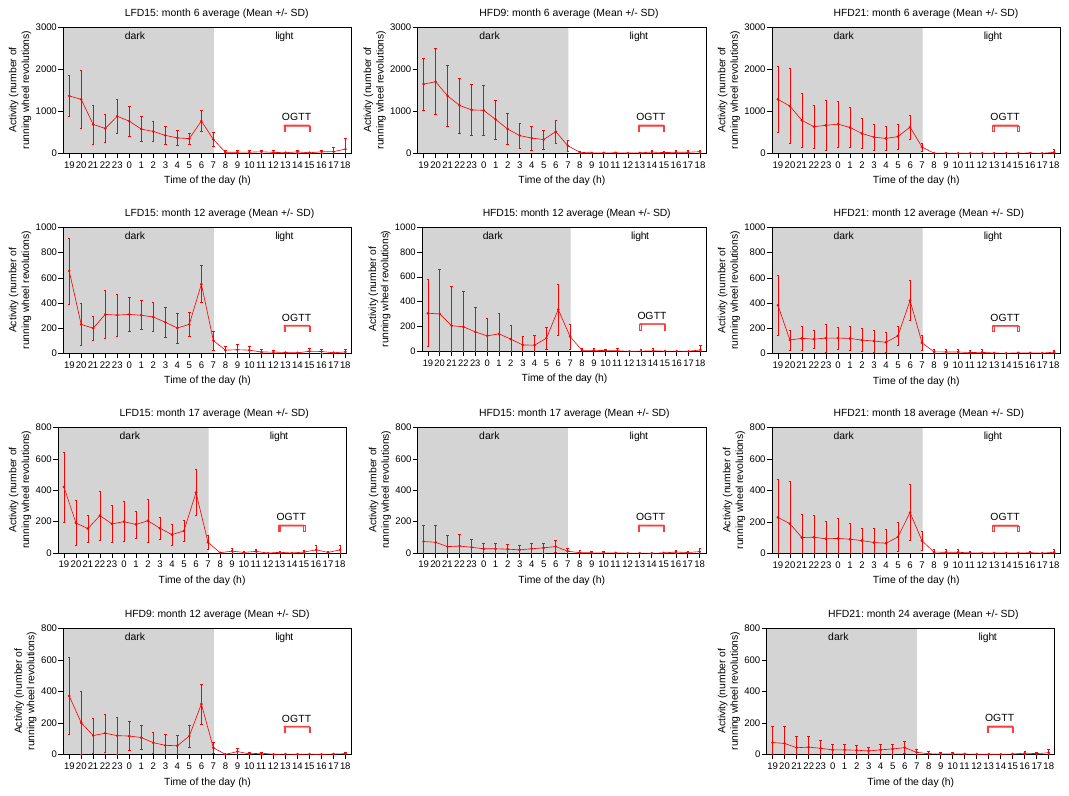
**Figure S1:** Average of running wheel activity in number of revolutions per hour. Data represent the average of all animals per experimental group on a given month (specified per panel). The diet group (LFD or HFD) is directly followed by a number, which indicates the experimental age group, as indicated in the materials and methods section. Dark and light phases are indicated, as well as the period of time in which the OGTT was conducted. Data are shown as mean ± SD, n = 16-26.

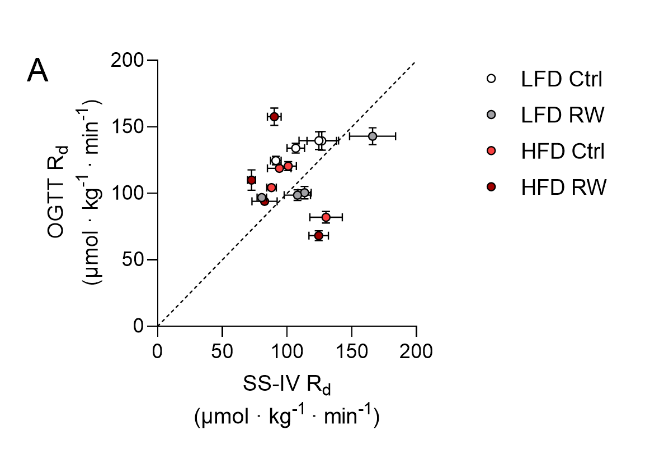

**Figure S2:** Comparison of the rate of glucose disposal (R_d_) at basal state between the tracer OGTT and steady-state intravenous infusion (IV-SS) data, age matched. The dashed line represents the identity line (y=x). Scatter plot of data plotted as mean ± SEM, n = 6-8.

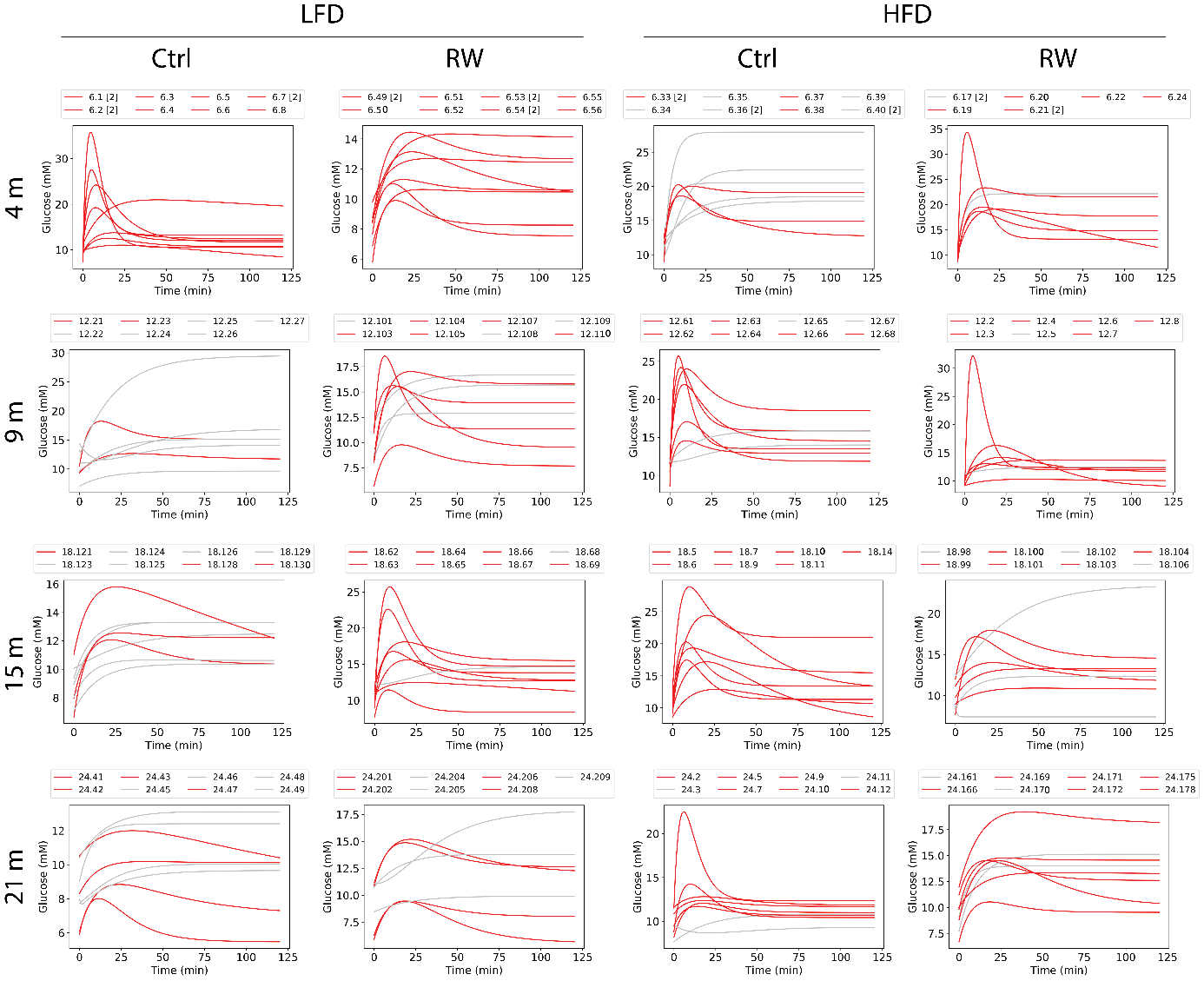

**Figure S3:** Individual fits for unlabeled glucose curves. Above each graph the mouse IDs are shown. Curves that showed an absorption followed by a clearance phase (red) were used for the EGP calculation. Curves in gray did not show the expected behavior and therefore were not included in the dataset to compute the EGP. This was done due to the high uncertainty in the obtained phenomenological constants for the unlabeled glucose in the absence of a clearance phase. In total, 37 animals were excluded, which were distributed among different groups.

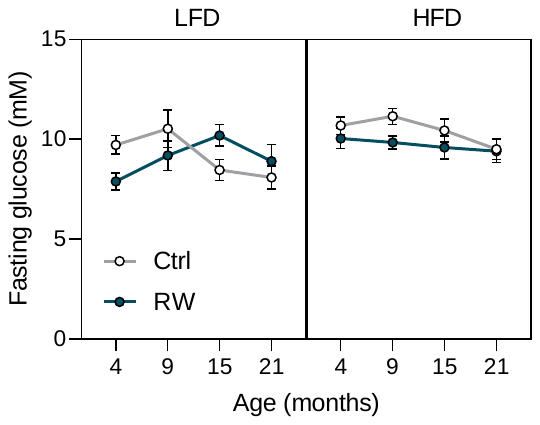

**Figure S4:** Fasting glucose values for each group. Data are shown as mean ± SEM, n = 6-8.

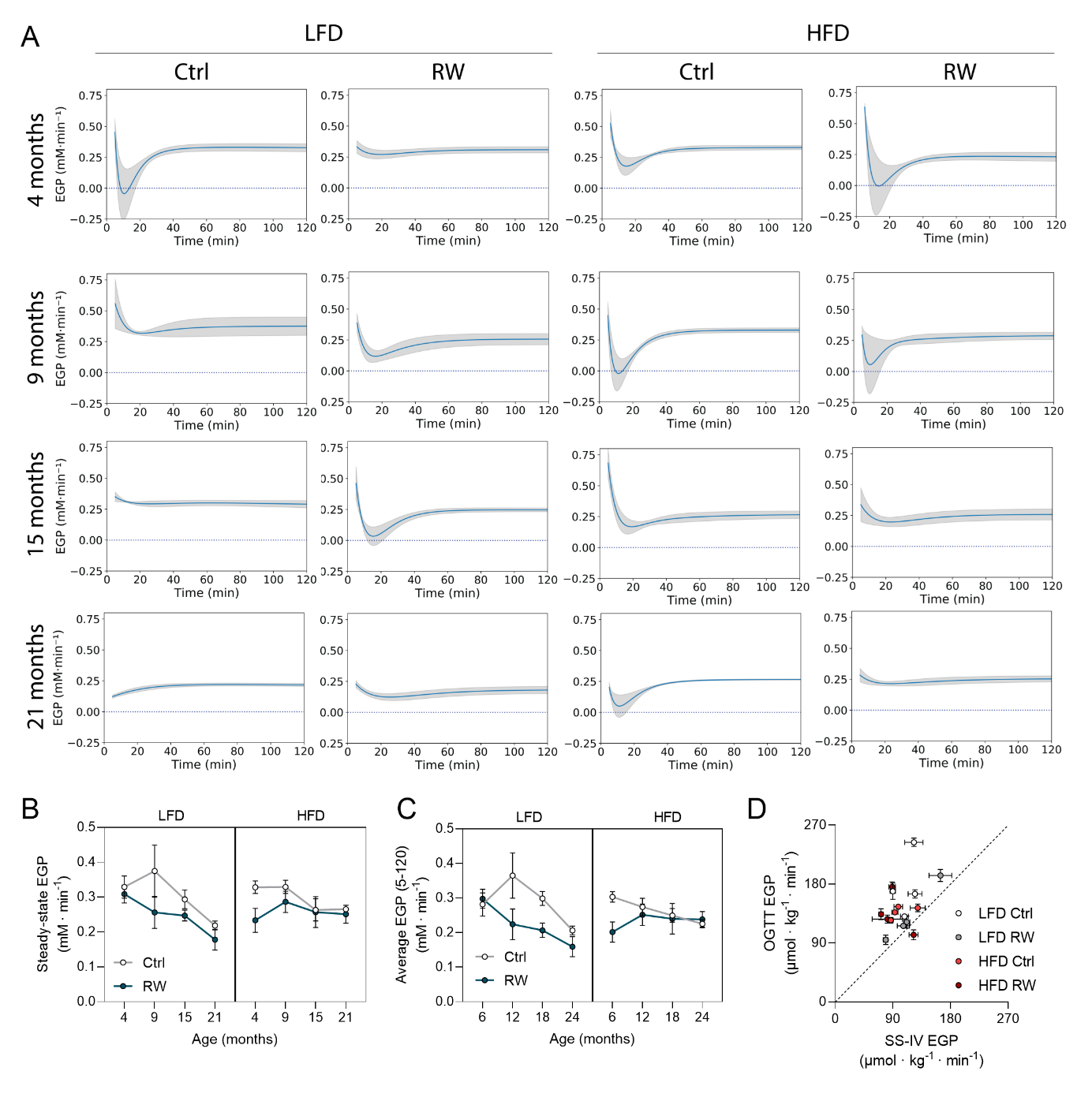

**Figure S5:** EGP* time courses in mM·min^-1^ (A). Each column represents a different diet and activity group, whereas each row represents a different age. LFD: low-fat diet, HFD: high-fat high-sucrose diet, Ctrl: sedentary mice, RW: mice submitted to voluntary running wheel. Mean EGP (line) ± SEM (shaded area) is shown per experimental group. Steady-state EGP* values in mM·min^-1^ (B) calculated from the curves (mean ± SEM). Average EGP* values in mM·min^-1^ (C) obtained from OGTT timeframe (5-120 min). Comparison of the steady-state EGP at basal state between the tracer OGTT and SS-IV datasets, age matched (D). The dashed line represents the identity line (y=x). Scatter plot of data plotted as mean ± SEM, n = 2-8.

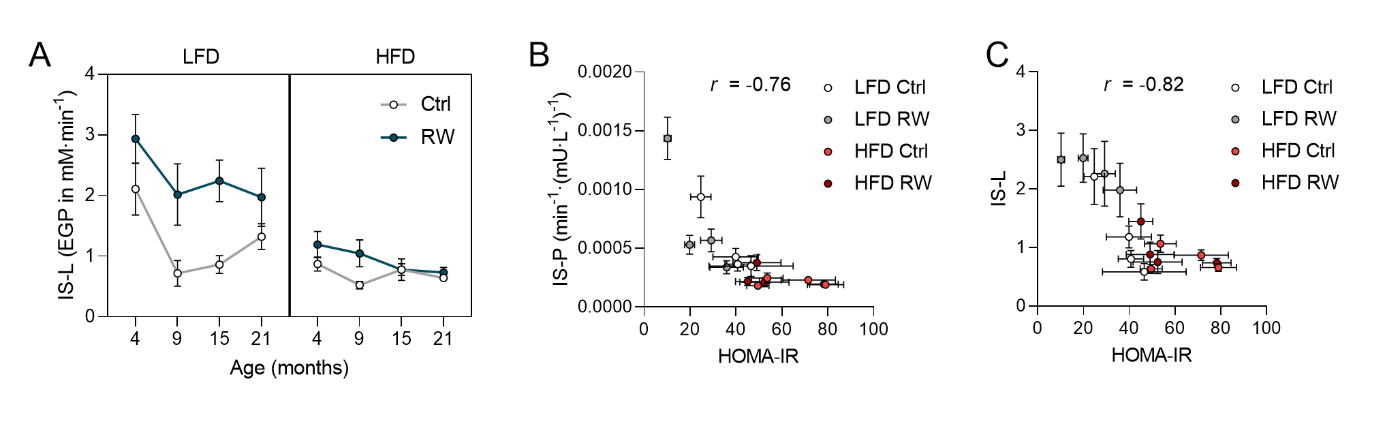

**Figure S6:** IS-L calculated from EGP in mM·min^-1^ (average from 5-120 minutes was used) (A). Data are shown as mean ± SEM, n = 2-8. Correlations between HOMA-IR and IS-P (B) or IS-L (C), Pearson correlation coefficients (*r)* are shown. Scatter plot of data plotted as mean ± SEM, n = 2-8.

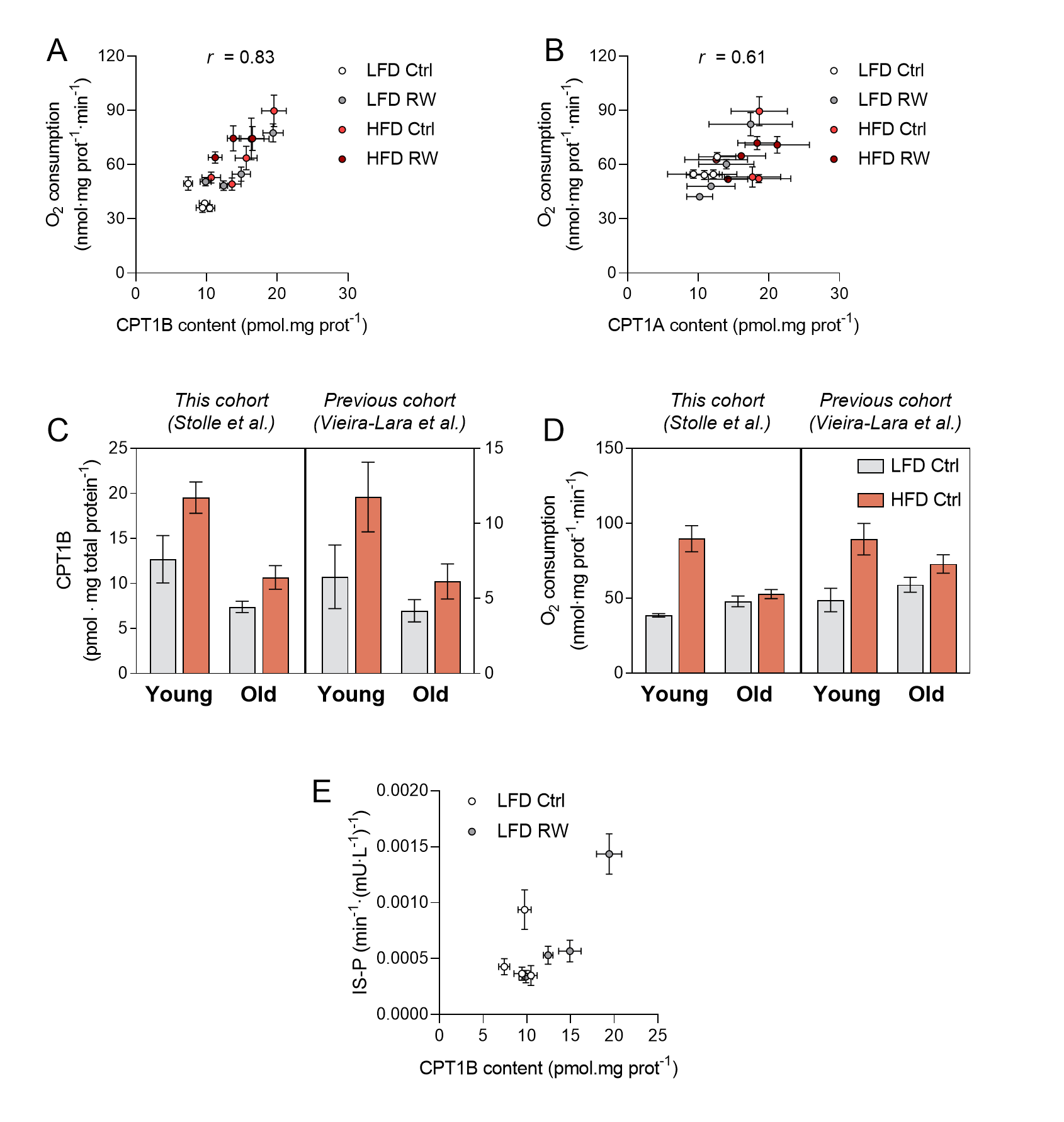

**Figure S7:** Correlation between CPT1B and oxidative capacity in the skeletal muscle, with the use of palmitoyl-CoA, carnitine and malate as substrates (A). Correlation between CPT1A and oxidative capacity in the liver, with the use of palmitoyl-CoA, carnitine and malate as substrates (B). Comparison between CPT1B levels (C) and oxidative capacity (palmitoyl-CoA, carnitine and malate as substrates) (D) in the skeletal muscle of mice from this present cohort (data from Stolle et al., 2018 (8), young = 4 months, old = 21 months, C57BL/6JOlaHsd background) and from a previous cohort (data from (9), young = 6 months, old = 21 months, C57BL/6J background). Correlation between CPT1B content and IS-P for LFD groups (E). Data are plotted as mean ± SEM, n = 4-8.
